# G-screen: Scalable Protein-Aware Virtual Screening through Flexible Ligand Alignment

**DOI:** 10.64898/2026.03.03.707320

**Authors:** Nuri Jung, Hahnbeom Park, Jinsol Yang, Chaok Seok

## Abstract

Virtual screening has long been a central computational tool for rational ligand discovery, enabling the systematic prioritization of candidate molecules from large chemical libraries. Although docking and related approaches that explicitly account for protein–ligand interactions have been developed and refined over several decades, achieving both reliable protein-aware interaction modeling and computational scalability remains an open challenge, particularly for ultra-large chemical spaces. Ligand-based methods are fast and robust but do not explicitly incorporate protein structure, whereas docking-based approaches model protein–ligand interactions more directly at substantially higher computational cost. Here, we present G-screen, a freely available and scalable protein-aware virtual screening framework designed for cases in which an experimentally determined or predicted reference protein–ligand complex structure is available. Rather than performing computationally intensive full docking with explicit pose sampling and optimization, G-screen rapidly generates alignment-guided pose hypotheses using a flexible global alignment algorithm (G-align). The resulting aligned poses are subsequently evaluated using protein-aware pharmacophore interactions derived from the reference complex, enabling explicit atomic-level interaction analysis while retaining the scalability and robustness of ligand-based alignment methods. Benchmarking on DUD-E, LIT-PCBA, and MUV datasets demonstrates that G-screen achieves competitive discrimination and early enrichment relative to representative ligand-based and docking-based methods, while maintaining millisecond-scale per-molecule runtimes under multi-threaded execution. These results position G-screen as a practical and scalable intermediate strategy between conventional ligand-based virtual screening and computationally intensive docking workflows for efficiently filtering ultra-large chemical libraries when a reference complex structure is available. G-screen and G-align are freely available at https://github.com/seoklab/gscreen and https://github.com/seoklab/galign, respectively.

**Scientific Contribution:** We have developed a scalable virtual screening framework for efficiently filtering ultra-large chemical libraries using a flexible global alignment algorithm combined with protein- aware pharmacophore evaluations and alignment-guided pose hypotheses. Despite explicitly capturing atomic-level interactions, the method remains highly efficient, maintaining millisecond-scale per-molecule runtimes under parallel execution. It achieves competitive discrimination and early enrichment, serving as an intermediate strategy between conventional ligand-based virtual screening and computationally intensive docking while combining the speed of ligand-based approaches with the structural context of traditional docking.

## 1. Introduction

Virtual screening (VS) has been a central computational strategy for ligand discovery against a given protein for several decades.^1–3^ It enables the prioritization of candidate molecules from large chemical libraries for experimental testing and remains a cornerstone of rational drug discovery.

VS methods broadly fall into two categories: ligand-based virtual screening (LBVS) and structure-based virtual screening (SBVS).^2, 3^ Ligand-based methods, including fingerprint similarity search^4^ and pharmacophore-based alignment,^5^ are among the simplest, fastest, and most robust approaches. By aligning candidate molecules to known ligands or extracting pharmacophore features from reference compounds,^3^ these methods indirectly account for protein–ligand interactions without explicitly modeling the protein. Their computational efficiency and robustness have made them enduring tools for large-scale screening.

To incorporate protein information more explicitly, structure-based approaches were developed, most prominently molecular docking.^6–8^ Docking predicts binding poses of ligands within a protein pocket and evaluates their compatibility. Although widely used, traditional docking remains computationally expensive for ultra-large libraries^9^ and can be limited by challenges in reliably predicting induced-fit conformational changes and binding geometry.^10–12^ Various strategies—including pre-filtering,^13^ active learning,^14, 15^ and fragment-based expansion^9^—have been proposed to mitigate cost, yet large-scale, accurate protein-aware screening remains challenging.

Recent advances in artificial intelligence have further reshaped the VS landscape. Deep learning– based pose prediction,^16–18^ AI-enabled docking pipelines,^19^ and joint structure prediction frameworks that simultaneously model protein folding and ligand binding (e.g., AlphaFold3^20^ and related approaches^21–23^) have demonstrated substantial progress in binding structure prediction under certain conditions. However, such methods are not yet routinely applicable to ultra-large library screening scenarios, where computational scalability and rapid throughput are critical.

Another emerging direction involves a protein-aware embedding approach, in which protein pockets and ligands are projected into shared representation spaces and interactions are modeled implicitly through learned similarity. A dense-retrieval framework^24^ has reported strong large-scale performance, highlighting the growing diversity of protein-aware screening strategies.

Meanwhile, the size of commercially accessible chemical libraries has expanded dramatically, reaching billions to trillions of compounds.^9, 25^ Screening such ultra-large libraries using a single high-cost method is computationally impractical. Efficient hierarchical workflows are therefore essential: rapid, scalable filtering must precede the application of more computationally intensive techniques. This creates a critical need for VS approaches that combine scalability with principled protein-aware interaction modeling.

In this context, we introduce G-screen, a scalable protein-aware virtual screening framework positioned between purely ligand-based alignment methods and full docking-based SBVS. G-screen assumes the availability of a reference protein–ligand complex structure, either experimentally determined or predicted. Rather than performing computationally intensive full docking with explicit pose sampling and optimization, G-screen rapidly generates alignment-guided pose hypotheses using a flexible global alignment algorithm (G-align) and evaluates protein-aware pharmacophore interactions derived from the reference complex. By combining fast structure alignment with explicit interaction analysis at atomic resolution, G-screen retains the scalability and robustness of ligand-based approaches while introducing principled protein-aware scoring without incurring the full computational cost of docking.

Comprehensive benchmarking across DUD-E,^26^ LIT-PCBA,^27^ and MUV^28^ demonstrates that G- screen achieves competitive performance relative to established ligand-based methods such as PharmaGist^29^ and docking-based methods such as AutoDock Vina.^30, 31^ Importantly, G-screen maintains millisecond-scale per-molecule runtimes under multi-threaded execution, enabling practical deployment for ultra-large screening campaigns as an initial filter for more computationally demanding virtual screening workflows. Collectively, these results position G-screen as a practical and scalable intermediate strategy between conventional ligand-based virtual screening and computationally intensive docking workflows when a reference complex structure is available.

## 2. Methods

### 2.1. G-screen method overview

G-screen is designed to identify molecules that interact with a target protein from a chemical library using a reference protein-ligand complex structure (**Figure 1**). The reference complex may be an experimentally determined structure or a predicted structure of a known or predicted bound ligand. The virtual screening pipeline begins by structurally aligning each library molecule to the reference complex. Based on this alignment, atomic interactions between the molecule and protein are mapped as protein-aware pharmacophore interactions. These interactions are then compared to those of the reference complex, with molecules ranked by interaction similarity, optionally incorporating shape similarity to the reference ligand.

**Figure 1.**
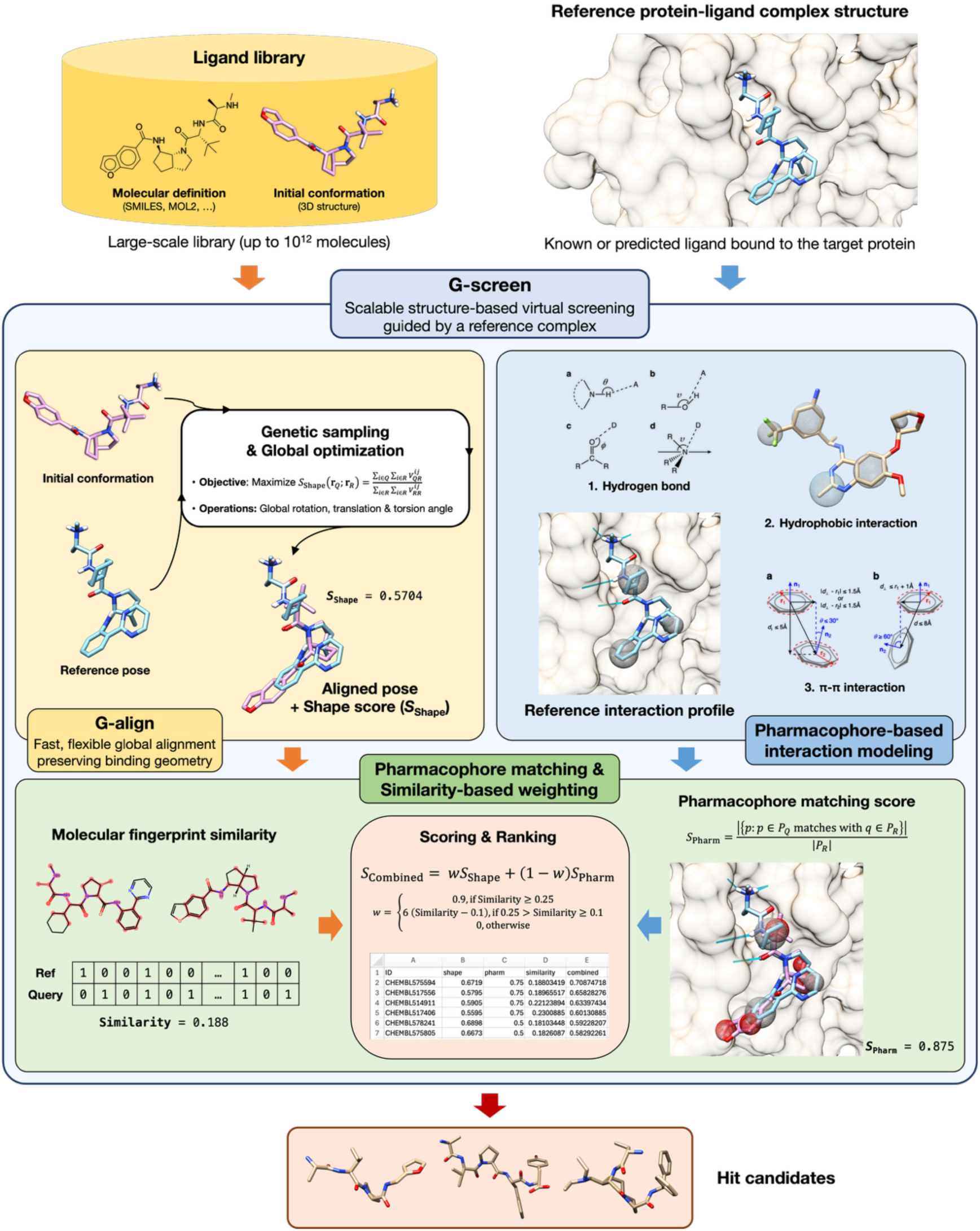
Overview of the G-screen virtual screening pipeline. G-screen performs scalable structure-based virtual screening guided by a reference protein–ligand complex. Library molecules are aligned to the reference ligand using the fast and flexible G-align algorithm (Section 2.2), preserving binding geometry, and are subsequently ranked by protein-aware pharmacophore interaction scoring (Section 2.3) at atomic resolution to identify hit molecules efficiently.

### 2.2. G-align: flexible structure alignment by global optimization

An important first step in evaluating the compatibility of a query molecule with the target protein pocket is G-align, which performs structure alignment of the query ligand to a fixed reference ligand. The goal is to identify hit molecules that can adopt chemical conformations compatible with the target protein pocket while maintaining a binding pose similar to that of the reference ligand. This flexible alignment is achieved by globally maximizing an objective function that measures shape similarity between the query and reference ligands over the translational, rotational, and torsional degrees of freedom of the query ligand. In the G-screen framework, the resulting aligned structures serve as alignment-guided pose hypotheses for subsequent protein-aware interaction evaluation, without requiring explicit docking-based pose sampling and optimization. Although a similar approach was previously implemented in the CSAlign algorithm,^12^ significant simplifications and algorithmic optimizations have been introduced here to enhance computational efficiency with minimal impact on alignment accuracy.

The G-align program was developed in early 2022 and has since been used in multiple in-house studies, including applications to chemical similarity search. For example, in CSearch,^32^ G-align was used as a proxy chemical similarity metric. However, CSearch provided only a brief description of the alignment procedure and scoring function, whereas the full formulation and algorithmic details (including the optimization procedure) were not reported. We therefore present the complete scoring function and algorithmic description of G-align here, which is intended to serve as the primary reference for the software.

#### 2.2.1. Shape score: objective for global optimization

The shape score, 𝑆_Shape_, measures the shape similarity between the query ligand structure (𝐫_Q_), defined by its translational, rotational, and torsional degrees of freedom, and the reference ligand structure (𝐫_𝑅_) as follows:

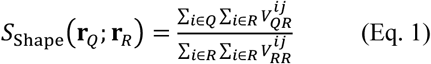

where 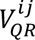 is the intersection volume between the *i*-th atom of the query ligand and the *j*-th atom of the reference ligand. Each atom’s radius is scaled down to 0.8 of its van der Waals radius to allow for local structural adjustments of both ligand and protein upon binding. The overlap volume is further scaled down by a factor of 0.7 if the SYBYL atom types differ, thereby penalizing chemical dissimilarity.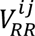 is defined analogously for reference ligand atoms.

This objective function is derived from the atomic match score of CSAlign,^12^ with the ligand internal energy term from GalaxyDock3^33^ omitted. Ligand internal energy, which reflects internal geometric and chemical strain, is instead implicitly accounted for by penalizing steric clashes during local optimization, as described in the following subsection (**2.2.2**).

#### 2.2.2. Global optimization algorithm

A genetic algorithm is used to evolve a fixed number of query ligand structures (10 in this study) through crossover and mutation to maximize the objective function, 𝑆_Shape_. The ligand structure that achieves the highest objective value is selected as the optimally aligned structure. Within the G-screen framework, this optimized aligned structure is interpreted as an alignment-guided pose hypothesis for subsequent protein-aware interaction scoring rather than as a fully docked binding pose. This approach is a modified version of CSAlign’s conformational space annealing algorithm,^12^ with the annealing component omitted.

In the genetic algorithm, ligand structures are represented as genes, which contain their internal coordinates including translational, rotational, and torsional degrees of freedom (DOFs). In this representation, a “crossover” occurs by mixing a subset of these DOFs between two genes, whereas “mutation” corresponds to a random change to a single DOF. The initial pool of structures (the “0th generation”) is generated from the input query ligand by randomly perturbing its DOFs, providing structural diversity for global optimization.

At each generation, 30 new trial structures are generated by crossover between structures selected from the current generation or between the current generation and the 0th generation, followed by mutation in the space of structural DOF. Each trial structure is created by crossover between a parent structure selected from the current generation with a probability proportional to its objective value and a partner structure chosen from either the current generation or the 0th generation, also weighted by objective value. This is followed by five mutation attempts on a randomly selected DOF, each with a probability of 50%.

The next generation of 10 structures is selected from the union of the current generation and the newly generated trial structures based on their objective values. This process is repeated until either the 50th generation is reached or until the maximum objective value remains unchanged for five consecutive generations.

All newly generated structures are locally optimized prior to objective evaluation using the Nelder-Mead algorithm^34^ at the gene level by maximizing a score that is defined as the shape score, 𝑆_Shape_, minus a steric clash penalty, 𝐸_Clash_. The clash penalty is calculated as the sum of pairwise clash contributions between all pairwise combinations of atoms in the query ligand, *Q*:

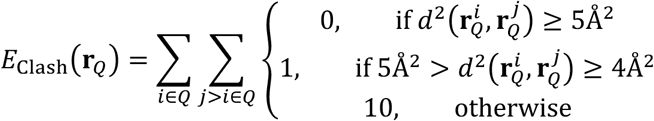

Where 𝑑^2^ denotes the squared Euclidean distance between atoms.

#### 2.2.3. Methods for benchmarking alignment methods

The performance of self-alignment, defined as aligning different structures of the same molecule, was evaluated using the same dataset previously employed to benchmark CSAlign.^12^ All three alignment methods—G-align, CSAlign, and Flexi-LS-align^35^—were executed with default parameters. The resulting aligned structures were assessed by calculating symmetry-corrected RMSD using the *obrms* program from the OpenBabel software suite (version 2.4.1).^36^

Time and memory usage for G-align, CSAlign, and Flexi-LS-align were measured on an Ubuntu 20.04 LTS system equipped with dual-socket AMD EPYC 7532 32-core processors at 2.4 GHz and 192 GB of RAM. Computational usage was monitored using the GNU time command (version 1.7).^37^

Because the aligned structures generated by G-align are subsequently used as alignment-guided pose hypotheses in the G-screen framework, preserving alignment accuracy while substantially improving computational efficiency is critical for scalable protein-aware virtual screening.

### 2.3. Assessment of Pharmacophore Interactions

The matching of pharmacophore interactions—hydrogen bonds, hydrophobic interactions, and π-π interactions (as defined below)—between the query complex 𝑃_Q_ and the reference complex 𝑃_𝑅_ is used to score the query ligand in its optimally aligned conformation (**Eq. 2**). This score serves as a coarse-grained measure of chemical interactions suitable for large-scale library screening. The score, 𝑆_Pharm_, can exceed 1 because pharmacophore features are evaluated independently in a pairwise manner between the query and reference complexes. As a result, multiple query features may contribute to the same reference feature when they are both geometrically compatible, and we selected this non-exclusive matching scheme to avoid imposing arbitrary assignment rules on this coarse-grained screening score.

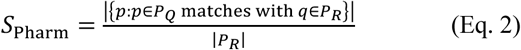

When evaluating 𝑆_Pharm_, all reference interactions are assigned once from the reference protein-ligand complex and subsequently treated as fixed during scoring for computational efficiency. Protein-side flexibility is not modeled, and no rotamer sampling or other protein conformational changes are considered, because the primary goal of G-screen is rapid and scalable large-scale virtual screening rather than detailed binding-pose refinement. Likewise, the aligned query ligand pose is scored as-is, without further conformational adjustment, in order to avoid the substantial computational cost associated with iterative docking-based optimization and to preserve the efficiency of the alignment-guided screening framework. The only exception is the limited rotameric conformation handling used for some hydrogen bond subtypes, as described in Section 2.3.1.

#### 2.3.1. Definition of hydrogen bond interactions

Hydrogen bonds between the aligned query ligand and the protein were detected based on hydrogen bond donor/acceptor types using modified criteria derived from a previous work by Mills and Dean,^38^ an algorithm also implemented in the UCSF Chimera.^39^

To simplify the procedure, a single distance and angle cutoff value was used for each hydrogen bond donor and acceptor type. The cutoff values were relaxed to allow for possible structural adjustments upon protein–ligand binding. The definitions of the angles are illustrated in **Figure 2**, and the modified criteria are summarized in **Table 1**. For all donor-acceptor pairs, the distance criterion was evaluated once per pair and was determined solely by the donor type, using either the hydrogen-acceptor atom or donor heavy atom-acceptor atom distance as specified in **Table 1**.

**Figure 2.**
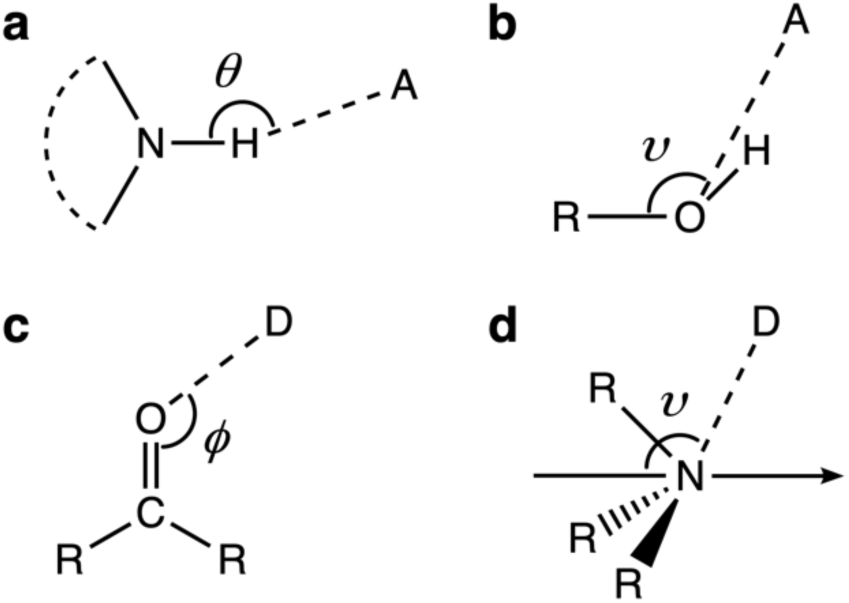
Illustration of the angles used for the hydrogen bond detection. A and D represent acceptor and donor heavy atoms, respectively. (a) *θ*; (b) donor *υ*; (c) *φ*; (d) acceptor *υ*.

**Table 1.** Hydrogen bond criteria by type.

| D/A<sup>a</sup> | Type | Definition | Cutoff values |  |
| --- | --- | --- | --- | --- |
|  |  |  | Distance (Å)<sup>b</sup> | Angle (°) |
| D | θ-τ | The hydrogen atom(s) of the donor atom are in-plane (e.g., $sp^2$-nitrogen atoms) | H-A<sup>c</sup>; 2.5 | $\theta > 135$ |
| D | ν-τ | The hydrogen atom(s) of the donor atom are out-of-plane (e.g., alcohols, primary and secondary amines) | D-A<sup>d</sup>; 3.5 | $150 > \nu > 90$ |
| A | ψ-φ | The lone pair(s) of the acceptor atom are in-plane (e.g., carbonyl groups, ethers, $sp^2$-nitrogen atoms) | (Implied from donor cutoff) | $\varphi \geq 90$ |
| A | ν-τ | The lone pair(s) of the acceptor atom are out-of-plane (e.g., alcohols, amines) | | $\nu \geq 90$ |

<sup>a</sup>D: Hydrogen bond donor, A: Hydrogen bond acceptor.
<sup>b</sup>Distance cutoffs were evaluated once per donor-acceptor pair and were determined solely by donor type.
<sup>c</sup>H-A, Hydrogen-acceptor atom distance.
<sup>d</sup>D-A, Donor-acceptor atom distance.

An additional improvement to the original algorithm was introduced by considering full rotameric conformations of terminal hydrogen bond donor and acceptor atoms. Candidate hydrogen bond donors and acceptors were identified using the *pybel* interface of the OpenBabel software.^36^

#### 2.3.2. Definition of hydrophobic interactions

Hydrophobic regions of a ligand, represented as spheres, were identified based on the method of Greene *et al.*,^40^ which provides the theoretical basis of the hydrophobe detection algorithm implemented in LigandScout.^5^

In brief, the original approach quantifies atomic hydrophobicity as ℎ = 𝑡𝑠, where 𝑡 is a topology-dependent hydrophobicity factor reflecting the local chemical environment, and 𝑠 is the solvent accessible surface area derived from the van der Waals radius and solvent exposure. A hydrophobic region is defined when the summed hydrophobicity exceeds a threshold of half the value of a methyl carbon, applied to small rings with fewer than eight heavy atoms or to clusters of connected hydrophobic atoms. The radius of each hydrophobic region is calculated as the average of distance of its constituent atoms from the region center each weighted by ℎ, scaled by the average 𝑡 value of the the region.

In the present implementation, two modifications were introduced. First, substituents containing fewer than three heavy atoms were considered as part of a ring when defining hydrophobic regions, to better represent the hydrophobic contribution of small substituents adjacent to aromatic or heteroaromatic systems. As a result, the hetero-ring is identified as a hydrophobic region in **Figure 3**b, in contrast to **Figure 3**a, reflecting the increased hydrophobic contribution from the methyl substituent. Second, hydrophobic groups consisting of fewer than three atoms that cannot be merged with neighboring hydrophobic regions are discarded. This effect is illustrated in **Figure 3**c, where the methyl group highlighted in **Figure 3**a and b is removed due to its inability to form a larger hydrophobic region.

**Figure 3.**
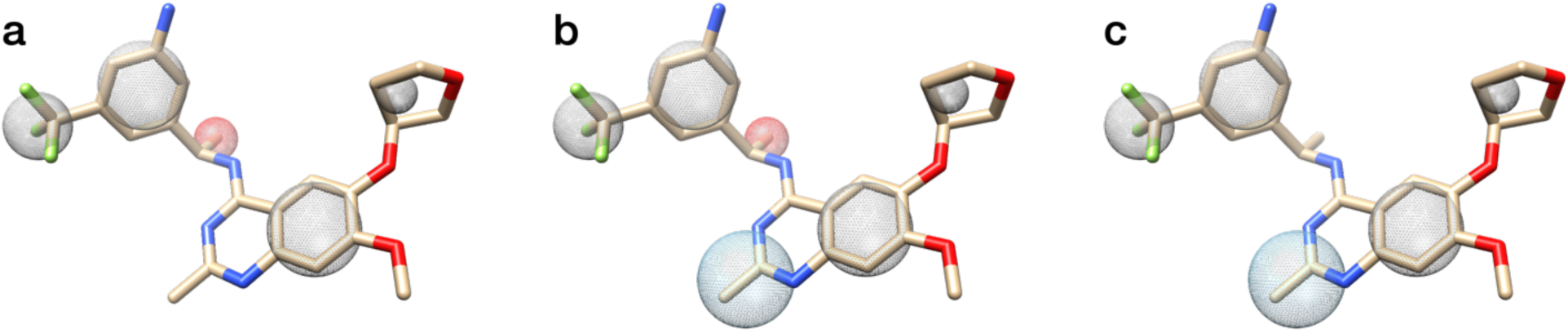
Effects of the modifications applied to the hydrophobic region detection algorithm. (a) The original algorithm. (b) Inclusion of small substituents connected to a ring, resulting in the identification of an additional hydrophobic region. (c) Removal of small hydrophobic groups after merging with neighboring hydrophobic regions. Added and removed hydrophobic regions are shown in sky-blue and red, respectively. The ligand example is taken from a PDB structure (PDB ID: 6SCM).

Unlike the other two pharmacophore interaction types, the hydrophobic region detection considers ligand atoms only, as a precise geometric definition of protein–ligand hydrophobic interactions remains ambiguous. Two hydrophobic regions are considered to match if the distance between their centers is either less than 1 Å or smaller than the sum of their radii.

#### 2.3.3. Definition of π-π interactions

Although π–π interactions can be considered a subtype of hydrophobic interactions, they are highly orientation-dependent unlike hydrophobic interactions, and often contribute significantly to binding stability and specificity and are therefore treated explicitly in this work. π-π interactions were detected following the criteria described by McGaughey *et al*.^41^ Two energetically favorable interaction subtypes were considered: parallel displaced and T-shaped interactions.

Aromatic rings were initially identified using the *pybel* interface of OpenBabel. To compensate for cases where aromaticity is incorrectly assigned, we additionally searched for 5–6 membered rings in which all ring atoms and directly connected neighbors are coplanar; these rings were also treated as aromatic. For fused, condensed, or polycyclic ring systems, ring candidates were enumerated by decomposing each ring system into its cycle basis, and each basis ring was treated as an independent aromatic unit if it satisfied the above criteria. Thus, fused systems were not represented by a single centroid or plane for the entire ring system; instead, each constituent ring was assigned its own centroid and normal vector.

π-π interaction detection was then performed at the level of individual ring pairs. First, the center of each aromatic ring was computed as the centroid of its atoms, and ring normal vectors were obtained from a best-fit plane. Ring pairs were subsequently evaluated against geometric criteria specific to each interaction subtype.

A parallel-displaced interaction was assigned when the angle between ring normal vectors was ≤ 30°, the inter-ring center distance was < 5Å, and at least one projected centroid offset was within 1.5 Å of the corresponding ring radius (**Figure 4**a).

**Figure 4.**
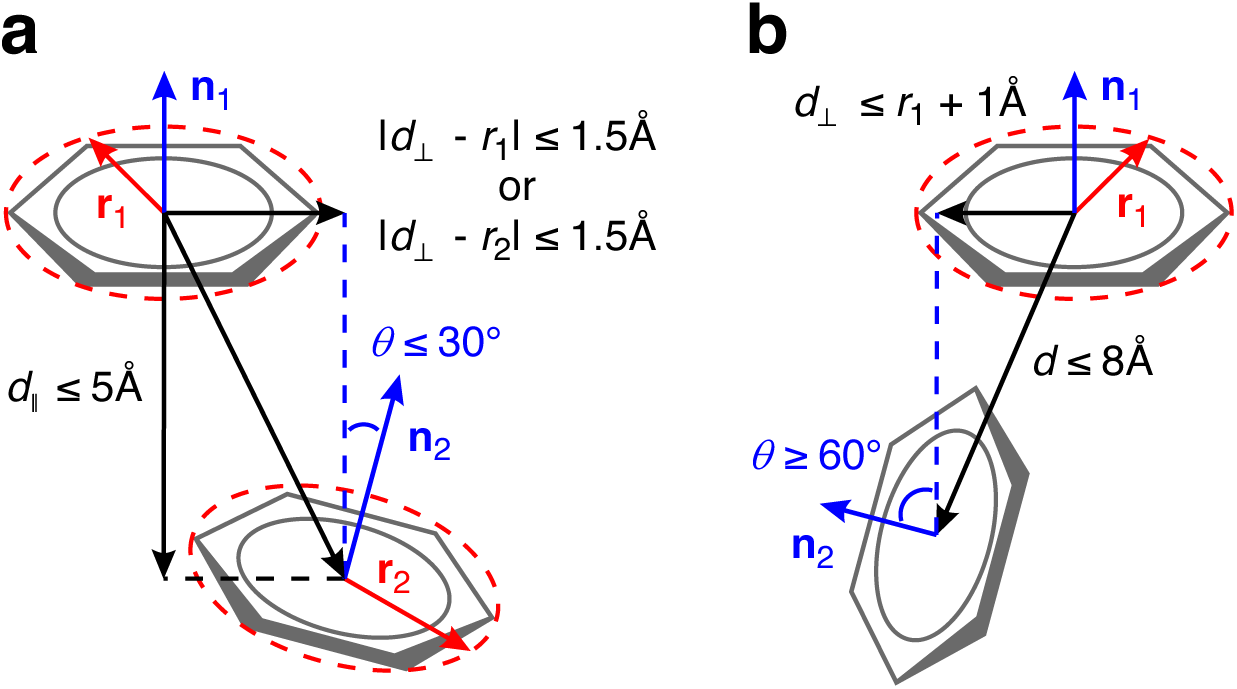
Illustration of the geometric criteria used to detect π–π interactions: (a) parallel-displaced and (b) T-shaped configurations.

A T-shaped interaction was assigned when the angle between the ring normal vectors exceeded 60°, the inter-ring center distance was < 8Å, and the parallel centroid offset was less than the radius of the base ring plus 1Å (**Figure 4**b). The base ring was defined as the ring with the smaller angle between its normal vector and the inter-center vector. When a fused system contributed multiple basis rings, each qualifying ring pair was evaluated independently.

#### 2.3.4. Ablation study of an explicit charged-interaction term

Charged interactions between positively and negatively ionizable groups are an important component of many protein-ligand recognition events.^42, 43^ In the main G-screen scoring scheme, charged interactions were not represented explicitly; instead, the proposed score was defined using hydrogen bonding, hydrophobic interactions, and π-π interactions. To assess whether explicit treatment of ionic interactions would improve screening performance, we additionally implemented charged interaction features following the LigandScout^5^ definition (**Method S1**).

This extended scoring variant was evaluated separately on the main benchmark set (Section 2.4.1) and compared with the default G-screen scoring scheme (**Table S1**). In our benchmarks, explicit inclusion of charged interaction features did not improve overall performance and often led to decreased performance for some targets. One possible reason is that charged interactions are strongly coupled to solvation and desolvation effects, such that favorable electrostatic interactions may be substantially offset by the energetic cost of desolvating charged groups. Accurately modeling this balance generally requires more detailed physical treatment than is practical within a coarse-grained, low-cost screening framework. These results suggest that, although charged interactions may be important for specific target classes, their explicit inclusion in the present formulation did not provide a consistent benefit across the benchmark sets. We therefore retained the original G-screen scoring formulation as the proposed method.

### 2.4. Virtual screening performance benchmark

#### 2.4.1. Benchmark datasets: DUD-E, LIT-PCBA, and MUV

To comprehensively evaluate the virtual screening performance of G-screen under varying levels of dataset bias and difficulty, we benchmarked the method on three widely used datasets: DUD-E, LIT-PCBA, and MUV.

The DUD-E dataset is one of the most commonly adopted benchmarks for virtual screening and provides a large number of targets with computationally generated decoys. While DUD-E enables systematic comparison across methods, its decoys are selected based on computational procedures, physicochemical property matching and similarity filtering,^26^ which may introduce bias favoring ligand similarity–based approaches.

To assess performance under more realistic and challenging conditions, we additionally evaluated G-screen on the LIT-PCBA and MUV datasets. Unlike DUD-E, these datasets are derived from experimentally measured high-throughput screening data,^27, 28^ in which decoys correspond to experimentally confirmed inactive compounds. As a result, actives and decoys in LIT-PCBA and MUV are substantially less distinguishable based on simple ligand similarity, making them more stringent benchmarks for protein-aware screening methods.

For each target in LIT-PCBA and MUV, a single reference protein–ligand complex structure was assigned for the main benchmark sets to maintain a consistent and fair evaluation protocol across methods. While some screening approaches can incorporate multiple reference ligands, others cannot, and aggregating scores across multiple references is not well defined for several workflows. We therefore selected one representative complex per target using predefined criteria, applied sequentially until a valid candidate was identified. Candidate structures for LIT-PCBA were taken from the benchmark dataset itself, whereas those for MUV were obtained from the Protein Data Bank (PDB).

The selection criteria were as follows: (1) the ligand is present in DrugBank; (2) experimental binding affinity data (IC50/EC50 or *K*i/*K*d) are reported; (3) when multiple candidates exist, the ligand with the strongest reported affinity is preferred; and (4) the protein is a known drug target of the ligand. If multiple structures satisfied these conditions, the complex included in the DUD-E dataset was preferred when available; otherwise, the complex with the fewest mutations and highest crystallographic resolution was selected. Targets lacking suitable experimental structures were excluded from the main benchmark sets.

To assess the sensitivity of G-screen to reference-template choice, we additionally constructed a cross-template validation subset from the remaining LIT-PCBA reference complexes and compared G- screen against the same baseline methods across multiple templates (**Method S2**). In parallel, to evaluate the applicability of G-screen to computationally predicted rather than experimentally determined templates, we generated AlphaFold3-predicted models for all selected reference complexes. These AF3- based reference models were used to examine performance changes associated with reference-structure substitution within the main benchmark targets. In addition, for the three MUV targets lacking experimental structures, AF3-predicted complexes were used in an exploratory benchmark to assess whether G-screen can be applied in the absence of known protein-ligand structures. For these targets, all methods were evaluated; for the main benchmark targets, AF3-based evaluation was performed only for G-screen to specifically assess the sensitivity of G-screen to predicted reference structures (**Method S3**).

To quantify dataset bias, we compared the average ECFP4 Tanimoto similarity between the reference ligand and active and decoy molecules across datasets (**Figure 5**). This analysis confirmed that DUD-E exhibits higher ligand similarity between actives and reference compounds, whereas LIT-PCBA and MUV present substantially lower similarity, thereby representing more challenging and less similarity-driven screening scenarios.

**Figure 5.**
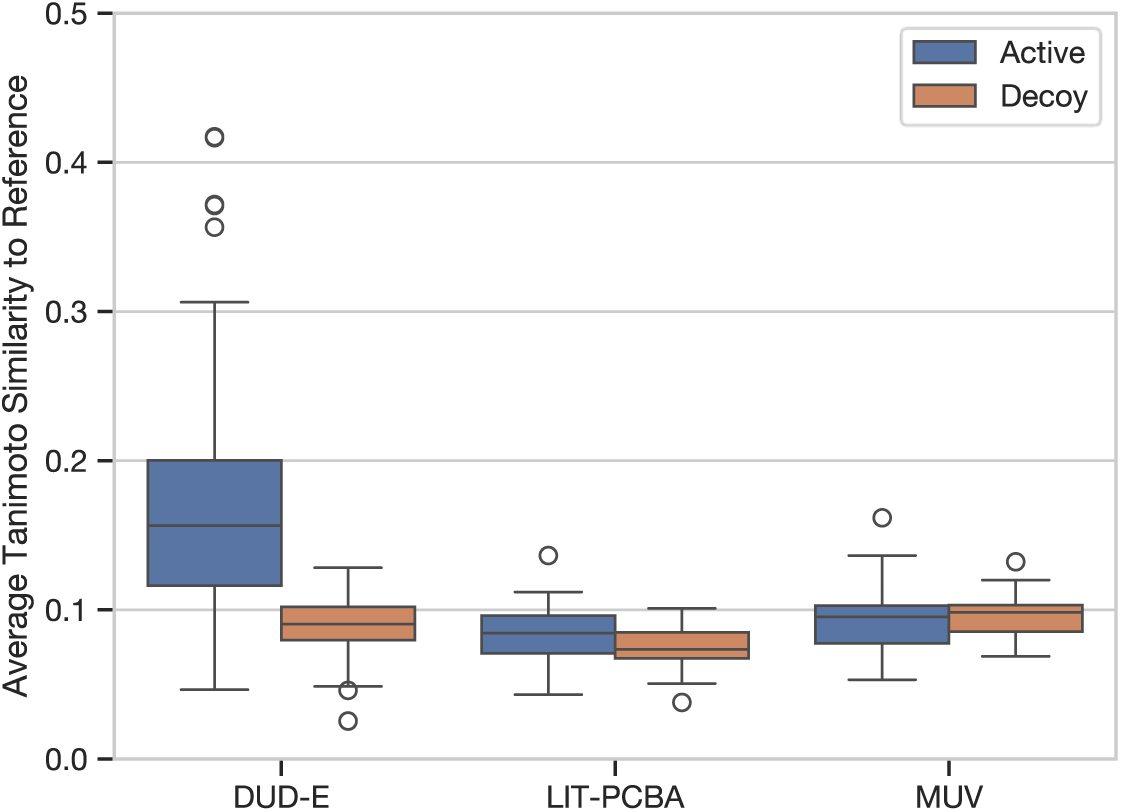
Active and decoy molecules in the LIT-PCBA and MUV set are less distinguishable based on ligand similarity than those in the DUD-E set, as reflected by the distributions of average ECFP4 Tanimoto similarity to the corresponding crystal ligand.

#### 2.4.2. Computational efficiency and parallel scalability

To assess the practical applicability of each screening workflow, we benchmarked computational cost in addition to screening accuracy. Because advances in multi-core CPU hardware make large-scale screening increasingly feasible, parallel scalability has become an important determinant of throughput, defined here as molecules screened per unit time. Single-threaded runtimes were used as the primary direct comparison across methods. Multi-threaded runtimes were additionally reported for methods with native support for parallel execution in order to illustrate practical scalability under realistic computational settings.

To evaluate computational efficiency, runtimes were measured on a randomly selected subset of 100 molecules per target, sampled without replacement from the combined active/decoy pool. Multi-threaded performance was additionally evaluated for methods that natively support parallel execution, namely G- align, G-screen, and AutoDock Vina. G-align and G-screen were benchmarked using 128 threads to assess parallel scalability. Because wall-clock timings at this level of parallelism can be noisy for small sample sizes, the 100-molecule subset was expanded to 5,000 evaluations by repeating the same 100 molecules 50 times with randomized ordering. AutoDock Vina was benchmarked in both single-threaded and multithreaded modes. Because performance gains saturated beyond 8 threads in our environment, multithreaded Vina timings were collected using 8 threads. Due to its substantially higher per-molecule cost, Vina runtimes were measured only on the original 100-molecule subset per target.

To assess the practical feasibility of methods that do not natively support multithreaded or multiprocess execution, namely Flexi-LS-align and PharmaGist, in a large-scale parallel screening context, we additionally analyzed empirical memory-scaling behavior as a function of the number of molecules processed per run for GS-S, GS-P/SP, Flexi-LS-align, and PharmaGist (**Method S4**). This supplementary analysis was used to evaluate whether comparable large-scale parallel execution settings were practically feasible across methods.

All resource-usage measurements were performed on Ubuntu 24.04 machines equipped with dual-socket AMD EPYC 7532 32-core processors at 2.4 GHz and 256 GB of RAM. Computational resource usage was monitored using the GNU time command (version 1.9).^37^

#### 2.4.3. Software and computational tools

Molecular graphics and analyses performed with UCSF Chimera, developed by the Resource for Biocomputing, Visualization, and Informatics at the University of California, San Francisco, with support from NIH P41-GM103311.^39^ Numerical data processing and visualization were performed with the help of seaborn,^44^ matplotlib,^45^ numpy,^46^ scipy,^47^ pandas,^48, 49^ scikit-learn,^50^ and jupyter notebook^51^ software packages. The NetworkX^52^ library was also used for the analysis of the 2D molecular graphs.

## 3. Results and Discussion

### 3.1. G-align: scalability with preserved accuracy

Before evaluating the overall virtual screening performance of G-screen, we first assess G-align, the flexible ligand alignment engine that enables the scalability of the G-screen pipeline. CSAlign^12^ has previously demonstrated high accuracy in flexible small-molecule alignment; however, its computational cost limits its direct applicability to large-scale screening. To address this limitation, we developed G- align as a simplified and multi-threaded variant designed to preserve alignment accuracy while substantially improving computational efficiency.

In the self-alignment benchmark (**Table 2**), G-align retained alignment accuracy comparable to that of CSAlign while achieving orders-of-magnitude improvements in computational efficiency. Under multi-threaded execution (128 threads), runtime was reduced by approximately four orders of magnitude relative to CSAlign, while maintaining stable memory usage.

**Table 2.**
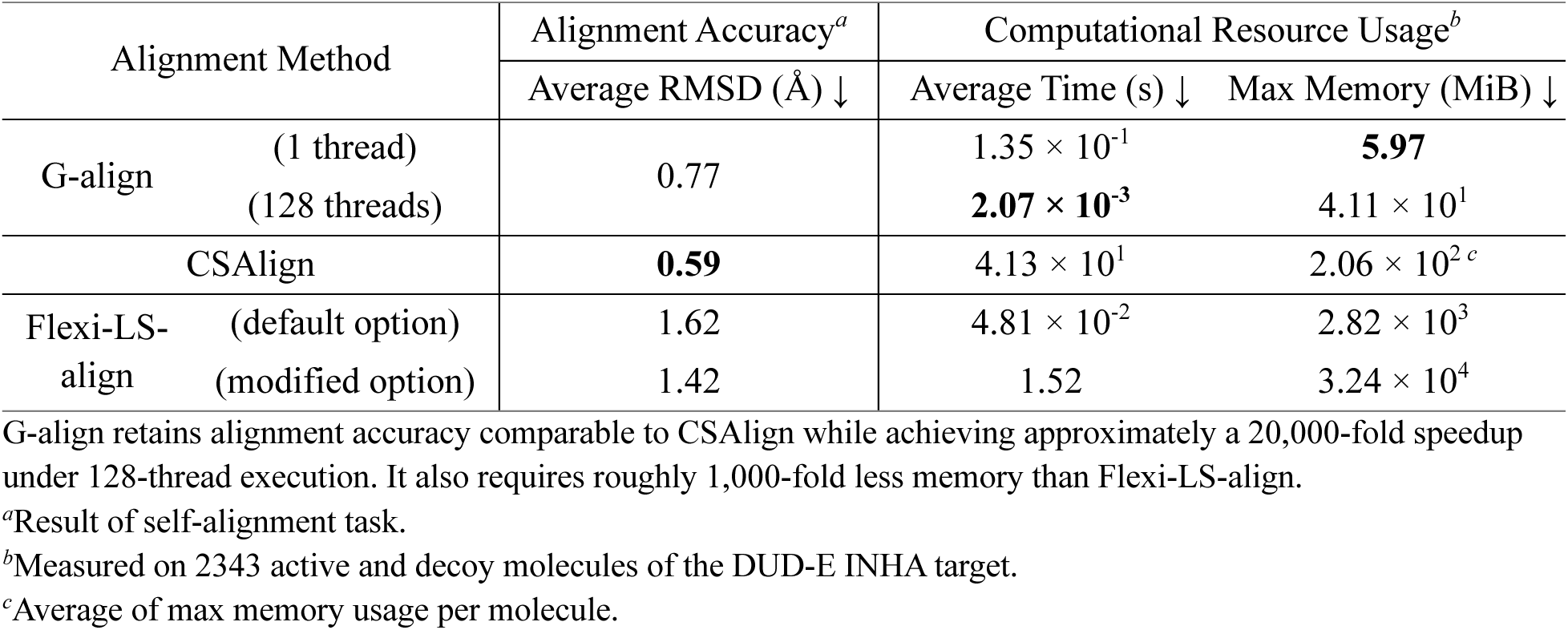
Performance and resource efficiency of G-align compared to existing flexible alignment methods.

We additionally compared G-align with Flexi-LS-align^35^. Although Flexi-LS-align is faster than the single-threaded version of G-align under default settings, it exhibits lower alignment accuracy, lacks native multi-thread support, and requires substantially higher memory (**Table 2**). In particular, its memory footprint increases considerably for large molecule sets, limiting its suitability for large-scale screening.

Taken together, these results demonstrate that G-align preserves the alignment accuracy of state-of-the-art flexible alignment methods while dramatically improving scalability through algorithmic simplification and parallel execution. This balance between accuracy and efficiency is critical for enabling the large-scale protein-aware screening strategy implemented in G-screen.

### 3.2. Scalable protein-aware virtual screening for ultra-large chemical libraries

We evaluated G-screen across the DUD-E, LIT-PCBA, and MUV benchmarks to assess complementary aspects of its applicability to ultra-large virtual screening, including screening performance across benchmark datasets (Section 3.2.1), computational scalability and resource efficiency (Section 3.2.2), practical deployment within hierarchical screening workflows (Section 3.2.3), and robustness to template selection and predicted reference structures (Section 3.2.4).

#### 3.2.1. Screening performance across DUD-E, LIT-PCBA, and MUV

To dissect the contributions of individual scoring components, we report results for G-screen using three different scoring schemes: the shape-only score (**Eq. 1**; GS-S), pharmacophore-only score (**Eq. 2**; GS-P), and their combination (GS-SP). Here, the combined score is defined as follows (**Eq. 3**):

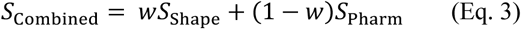

where 𝑤 is the weight factor for the shape compatibility score. The weight is determined from the ECFP4 fingerprint Tanimoto coefficient (Similarity) of the query molecule and the reference ligand.

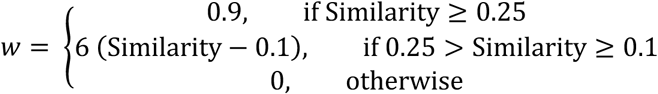

This dynamic weighting scheme drives the combined score to focus more on pharmacophore similarity as the query becomes dissimilar to the reference.

To analyze the performance of the G-screen methods alone, we compared the G-screen variants (GS-S, GS-P, and GS-SP) against Flexi-LS-align (ligand alignment method),^35^ PharmaGist (ligand-based pharmacophore method),^29^ and AutoDock Vina (structure-based docking method).^30, 31^ Performance was assessed using AUROC and early enrichment factors computed for the top 0.1%, 1%, and 5% of ranked molecules (EF0.1%, EF1%, and EF5%) for each method. Because EF1% provides a practical and statistically stable measure of early enrichment across benchmark targets, the following discussion primarily focuses on EF1%. This was evaluated together with similarity enrichment to quantify potential ligand similarity bias (**Table 3**; **Figure S1**). Similarity enrichment was defined as the ratio of the average ECFP4 Tanimoto similarity of the top 1% of ranked molecules to that of all actives and decoys for a given target.

**Table 3.** Virtual screening performance on the three benchmark datasets.

| Dataset | Screening Method | Screening Performance (↑) | | | | Similarity Enrichment<sup>a</sup> (Top 1%) | Relative Avg. Runtime<sup>b</sup> (↓) | |
| --- | --- | --- | --- | --- | --- | --- | --- | --- |
| | | AUROC | EF0.1% | EF1% | EF5% | | Serial | Parallel<sup>c</sup> ($N_{\max}$) |
| DUD-E | GS-S | <u>0.75</u> | <u>38.15</u> | <u>19.71</u> | **7.03** | +86.7% | 0.87x | 0.014x (128) |
|  | GS-P | 0.72 | 21.70 | 13.62 | 5.86 | +54.3% | 1x | 0.023x (128) |
|  | GS-SP | **0.76** | 15.02 | 14.38 | 6.92 | +57.3% |  |  |
|  | Flexi-LS-align | 0.74 | **43.19** | **20.87** | **7.03** | +104.6% | 0.37x | N/A (memory scaling) |
|  | PharmaGist | 0.73 | 35.73 | 18.78 | 6.84 | +86.5% | 4.0x | N/A (memory scaling) |
|  | AutoDock Vina | 0.70 | 14.95 | 8.49 | 4.32 | +24.6% | 513x | 105x (8) |
| LIT-PCBA | GS-S | 0.56 | 11.58 | 3.67 | 1.88 | +29.4% | 0.84x | 0.013x (128) |
|  | GS-P | 0.56 | 6.46 | 3.78 | 2.03 | +14.8% | 1x | 0.025x (128) |
|  | GS-SP | 0.56 | <u>15.73</u> | <u>4.80</u> | <u>2.20</u> | +22.7% |  |  |
|  | Flexi-LS-align | <u>0.57</u> | 15.29 | 3.31 | 1.77 | +40.6% | 0.32x | N/A (memory scaling) |
|  | PharmaGist | **0.59** | **21.63** | **5.54** | **2.78** | +29.8% | 27x | N/A (memory scaling) |
|  | AutoDock Vina | 0.53 | 3.28 | 2.25 | 1.50 | +12.2% | 425x | 91x (8) |
| MUV | GS-S | 0.47 | 2.62 | 0.75 | 1.10 | +1.9% | 0.84x | 0.013x (128) |
|  | GS-P | <u>0.56</u> | 1.61 | 1.94 | <u>1.94</u> | +2.9% | 1x | 0.024x (128) |
|  | GS-SP | <u>0.56</u> | **2.87** | **2.43** | **2.21** | +2.2% |  |  |
|  | Flexi-LS-align | 0.49 | 0.00 | 0.73 | 0.73 | +28.0% | 0.42x | N/A (memory scaling) |
|  | PharmaGist<sup>d</sup> | **0.58** | 0.00 | 1.76 | 1.86 | +14.6% | 2.1x | N/A (memory scaling) |
|  | AutoDock Vina | 0.53 | <u>2.62</u> | <u>2.27</u> | 1.14 | +2.8% | 413x | 87x (8) |

<sup>a</sup>Similarity enrichment was defined as the ratio of the average ECFP4 Tanimoto similarity of the top 1% ranked molecules to that of all actives and decoys for a given target.
<sup>b</sup>Computational resource usage was measured separately using 100 randomly selected molecules per target. For G-screen methods, each 128-thread run used the same molecules repeated 50 times in shuffled order to reduce parallelization noise. Values are reported relative to single-thread GS-SP run.
<sup>c</sup>Parallel computation times are reported at the highest effective parallelism level for each benchmarked method (up to 128 threads). Methods exhibiting approximately linear memory scaling with database size were not included in the parallel benchmark, because memory usage was anticipated to limit practical scaling more strongly than thread count; these cases are therefore labeled as “N/A (memory scaling).”
<sup>d</sup>PharmaGist failed on two targets, MUV-712 and MUV-846, due to excessive memory usage and an unsupported boron-containing reference ligand, respectively.

On the DUD-E dataset (average 14,059 ligands across 102 targets; **Table 3**), GS-SP achieved an AUROC of 0.76, comparable to Flexi-LS-align (0.74) and exceeding AutoDock Vina (0.70). Early enrichment at EF1% reached 14.38, close to Flexi-LS-align (20.87) and PharmaGist (18.78). As expected for DUD-E, where actives tend to exhibit higher similarity to the reference ligand, ligand-based alignment performs strongly. Nevertheless, GS-SP exhibited lower similarity enrichment (+57.3%) than Flexi-LS-align (+104.4%), suggesting reduced reliance on simple ligand similarity.

The LIT-PCBA dataset (average 176,798 ligands across 15 targets) represents a more stringent benchmark with substantially reduced ligand similarity bias (**Table 3**). Across methods, AUROC values were modest (0.53–0.59), reflecting intrinsic dataset difficulty. GS-SP achieved AUROC 0.56, comparable to Flexi-LS-align (0.57) and close to PharmaGist (0.59), while improving EF1% (4.80) relative to GS-S (3.67) and GS-P (3.78). Similarity enrichment remained moderate (+22.7%), indicating that performance was not primarily driven by similarity to the reference ligand.

The MUV dataset (average 14,698 ligands across 14 targets) constitutes the most stringent evaluation, with minimal ligand similarity bias (**Table 3**). In this setting, purely shape-based alignment (GS-S) yielded AUROC 0.47, whereas protein-aware scoring significantly improved performance: GS-P and GS- SP both achieved AUROC 0.56. At EF1%, GS-SP reached 2.43, outperforming GS-S (0.75) and Flexi-LS-align (0.73), and comparable to AutoDock Vina (2.27). These results demonstrate that explicit protein-aware pharmacophore scoring becomes increasingly beneficial as ligand similarity bias diminishes.

Across all three benchmark sets, GS-SP consistently balanced discrimination performance and similarity enrichment, maintaining competitive AUROC while avoiding excessive dependence on ligand similarity. Collectively, these results indicate that G-screen achieves robust protein-aware screening performance across datasets with varying similarity regimes.

#### 3.2.2. Computational scalability and resource efficiency

We further evaluated the computational efficiency of all methods across the DUD-E, LIT-PCBA, and MUV benchmarks (**Table 3** and **Figure 6**). Single-threaded measurements were used as the primary matched comparison across methods. Under this setting, GS-SP remained among the faster methods tested despite explicitly incorporating protein-aware interaction scoring, with per-molecule runtime comparable to, and in some cases lower than, those of ligand-based methods such as Flexi-LS-align and PharmaGist. To illustrate implementation scalability, G-screen variants were additionally benchmarked under 128-thread execution, under which GS-SP ranked second only to GS-S, the shape-only component of the GS-SP pipeline.

**Figure 6.**
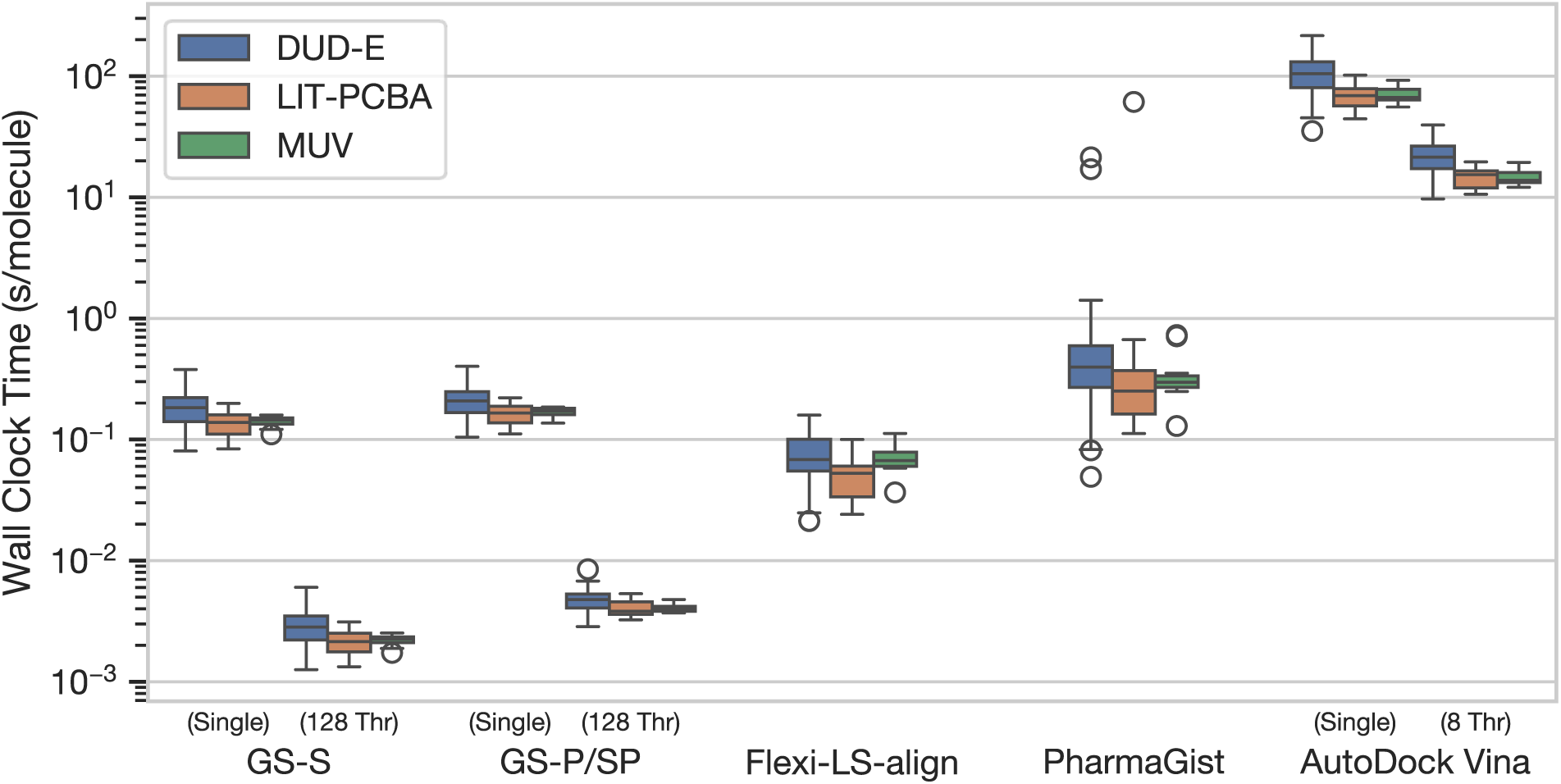
Per-molecule computational cost of the benchmarked methods, reported as elapsed wall-clock time (s/molecule; logarithmic scale). Single-threaded runtimes are shown for all methods as the primary matched comparison. G-screen variants are additionally shown in 128-thread mode to illustrate implementation scalability; GS-P and GS-SP are combined because their runtimes were similar. AutoDock Vina is shown in single-thread and 8- thread modes, as speedup saturated beyond 8 threads in our benchmark environment. Under parallel execution, GS- S and GS-P/SP achieve millisecond-scale runtimes, whereas the only other protein-aware method considered, AutoDock Vina, requires 2–3 orders of magnitude longer per-molecule runtime.

PharmaGist showed competitive screening performance on several benchmarks, but its computational requirements were highly target-dependent. It could not be evaluated on two MUV targets (MUV-712 and MUV-846) because of excessive memory usage and an unsupported boron-containing reference ligand, respectively. In the LIT-PCBA and MUV datasets, PharmaGist required substantial memory allocations, reaching up to approximately 180 GB for 500–1,000 candidate molecules, and for MUV-712 it exceeded 192 GB of RAM even for 10 molecules. These observations indicate that memory usage should be considered when applying this type of ligand-based pharmacophore method in large-scale screening settings. Consistent with this observation, separate memory-scaling analysis of Flexi-LS-align and PharmaGist confirmed that these memory limitations were substantial in practice: under a 256 GB RAM budget, approximately 30% of benchmarked targets were predicted to become infeasible at 128-thread execution when the workload reached about 1,000 molecules per core (**Figure S2**). This practical limitation is reflected in **Table 3** as “N/A (memory scaling)” for the corresponding parallel- runtime entries.

AutoDock Vina, the other protein-aware method included in the benchmark, required 2–3 orders of magnitude longer runtime per molecule than GS-S and GS-P/SP even under parallel execution. In addition, Vina speedup saturated at approximately 8 threads in our benchmark environment, limiting its scalability for large-scale virtual screening.

Taken together, these results demonstrate that G-screen offers a favorable balance between screening accuracy and computational efficiency. Combined with its free availability and efficient multithreaded scalability, G-screen provides a practical and accessible solution for large-scale protein-aware virtual screening, particularly in scenarios where docking-based approaches are computationally prohibitive.

#### 3.2.3. Hierarchical screening workflow for ultra-large virtual screening

G-screen is designed to serve as a rapid pre-filtering method that can be applied upstream of more computationally demanding virtual screening approaches, owing to its computational efficiency and scalability. To evaluate this use case, we constructed a hierarchical screening workflow in which molecules with identical GS-SP scores in the early enrichment region were further ranked using two established methods that performed strongly in our benchmark comparison: PharmaGist and AutoDock Vina. Enrichment was then evaluated using the enrichment factor among the top 1% of ranked compounds (EF1%; **Figure 7**).

**Figure 7.**
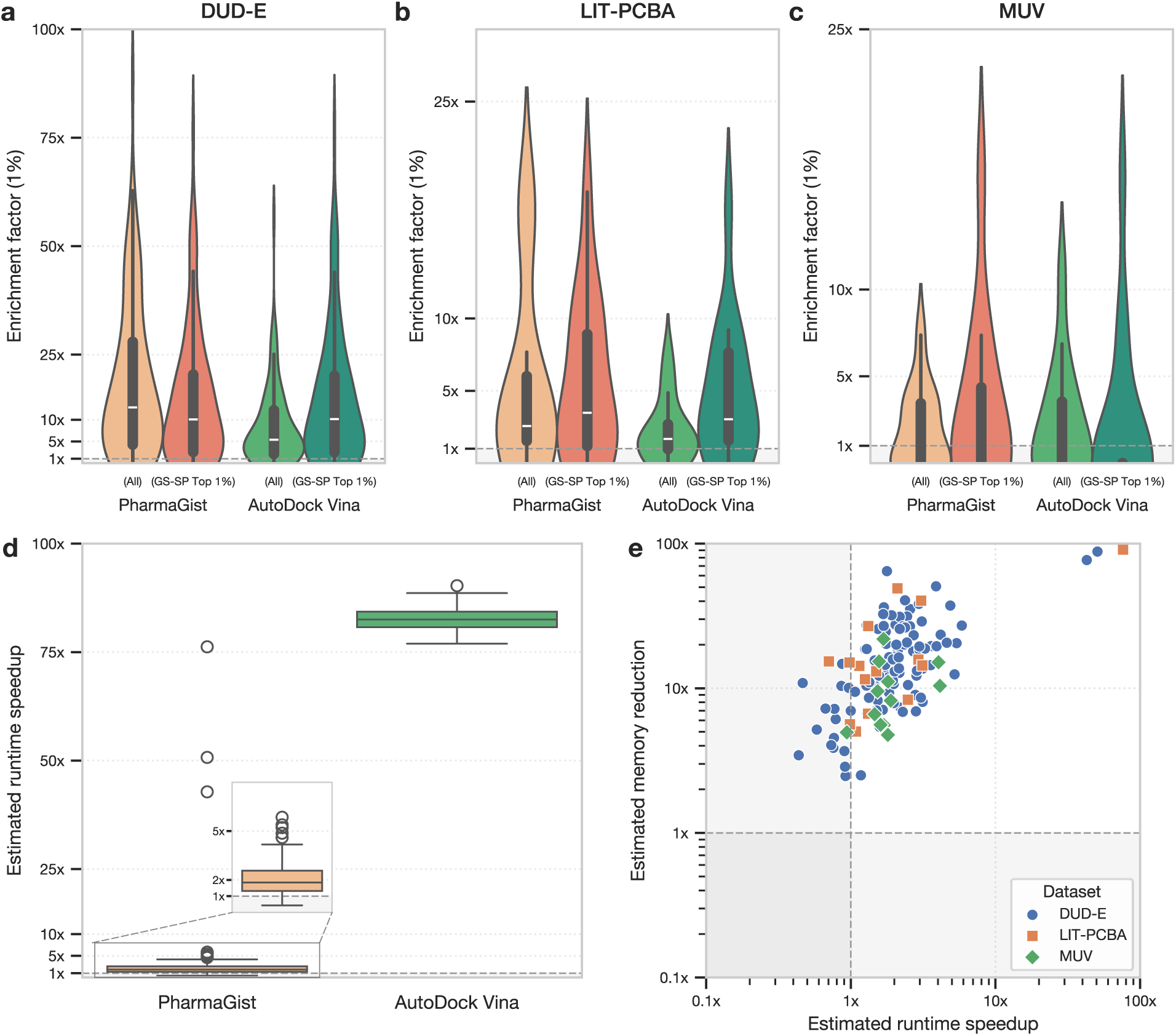
GS-SP pre-filtering improves the efficiency of downstream virtual screening while maintaining early enrichment. **(a-c)** Enrichment factor at 1% (EF1%) evaluated on the **(a)** DUD-E, **(b)** LIT-PCBA, and **(c)** MUV datasets. Violin plots compare PharmaGist and AutoDock Vina applied to the full compound library (“All”) with a hierarchical GS-SP pipeline (“GS-SP Top 1%”), in which GS-SP is used as the primary ranking method and the baseline method is applied only as a secondary score to resolve GS-SP score ties within the top 1% enrichment region. **(d)** Estimated runtime speedup of the GS-SP hierarchical pipeline relative to applying each baseline method to the full compound library. The inset shows a magnified view of the speedup distribution for PharmaGist. **(e)** Log-log scatter plot of estimated memory reduction versus estimated runtime speedup for PharmaGist across the evaluated datasets. Each point represents one target, colored by dataset.

We compared each baseline method applied to the full compound library (“All”) with the corresponding tie-breaking workflow, in which the baseline method was applied only to molecules tied by GS-SP within the top 1% region (“GS-SP Top 1%”). The estimated runtime speedup was calculated from the single-threaded average runtime per molecule for each method summarized in **Table 3** as:

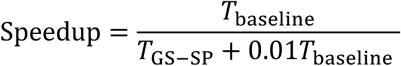

Where 𝑇_baseline_ and 𝑇_GS–SP_denote the average per-molecule runtimes of the downstream baseline method and GS-SP, respectively. Memory reduction for PharmaGist was estimated analogously from the memory-scaling analysis (**Method S4** and **Figure S2**), using the log-log least-squares fitted relationship over the measured range to estimate memory requirements for the full candidate set per target. Because the downstream method is applied only within the top 1% GS-SP region, both runtime speedup and memory reduction have a theoretical upper bound of 100-fold.

Across DUD-E, LIT-PCBA, and MUV, GS-SP-based pre-filtering followed by baseline tiebreaking generally preserved or improved EF1% relative to applying the baseline methods to the full library (**Figure 7**a-c). Notably, both downstream methods showed modest EF1% gains after GS-SP tiebreaking on the LIT-PCBA dataset. These results indicate that GS-SP can substantially reduce the candidate set requiring downstream evaluation without systematically compromising early enrichment.

The computational benefit of this tie-breaking workflow was substantial, particularly for AutoDock Vina (**Figure 7**d). Because only molecules in the top 1% GS-SP region are passed to the downstream method, the estimated runtime reduction was large for docking-based screening, yielding a median speedup of approximately 80-fold and approaching the theoretical upper bound of 100-fold. PharmaGist showed a more modest speedup, consistent with its different computational profile as a shape- and pharmacophore-based screening method. For PharmaGist, GS-SP filtering also consistently reduced estimated memory requirements (**Figure 7**e), with most targets showing simultaneous runtime and memory reductions. Together, these results support GS-SP as a scalable and complementary pre-screening method that rapidly prioritizes molecules while allowing more computationally expensive methods to resolve local ranking ambiguity among compounds tied by the GS-SP scoring scheme.

#### 3.2.4. Robustness to template selection and predicted structures

Although the main benchmark used a single reference complex per target for computational tractability, additional evaluation on a separate cross-template validation subset showed that the overall performance trend was broadly preserved across alternative templates. In this analysis, paired Wilcoxon signed-rank tests showed that GS-SP achieved significantly higher AUROC than Flexi-LS-align and AutoDock Vina after Benjamini-Hochberg false discovery rate correction (*p* < 0.01). As expected for a template-based method, G-screen performance remained dependent on template choice in both AUROC and EF1%. However, paired Wilcoxon signed-rank tests on cross-template metric dispersion did not detect statistically significant differences between G-screen and any of the benchmarked baselines after the same multiple-testing correction (*p* > 0.05; **Table S2, Figure S3**).

To further assess robustness to reference-structure quality, we evaluated G-screen on the main benchmark targets using AF3-predicted reference structures in place of experimentally determined complexes (**Table S1**). As expected for a method that evaluates protein-ligand interactions at atomic resolution, performance on modeled structures was generally lower than that obtained with experimental references. However, the decrease was modest, and G-screen remained competitive with the benchmarked baseline methods. For example, in the LIT-PCBA dataset, EF0.1% remained essentially unchanged (15.72 using AF3 models versus 15.73 using experimental structures), while EF1% was slightly improved (5.08 versus 4.80).

We also performed an exploratory benchmark on three MUV targets lacking experimental structures, in which GS-SP achieved an AUROC of 0.52 and an EF1% of 2.29 (**Table S3**). In this setting, GS-SP remained competitive with the best-performing baseline, AutoDock Vina, which achieved an AUROC of 0.52 and an EF1% of 3.46. Taken together, these results suggest that G-screen can remain practically useful even when only computationally predicted reference structures are available.

### 3.3. Molecular interpretation and applicability domain of G-screen

#### 3.3.1. Molecular interaction-based interpretation of screening behavior

To better understand the behavioral differences between protein-aware and ligand-based screening approaches, we analyzed representative targets at the molecular interaction level. Rather than focusing solely on ranking metrics, we examined how each method prioritizes molecules in terms of interaction patterns relative to the reference protein-ligand complex.

First, to evaluate the contribution of ligand similarity to ranking behavior, we compared ECFP4 Tanimoto similarity enrichment for the top 1% ranked molecules (**Figure 8**). Across most targets, G- screen exhibited lower similarity enrichment than PharmaGist, indicating a reduced dependence on ligand similarity during prioritization. This observation supports the interpretation that protein-aware scoring promotes the selection of structurally diverse active molecules while maintaining competitive screening performance.

**Figure 8.**
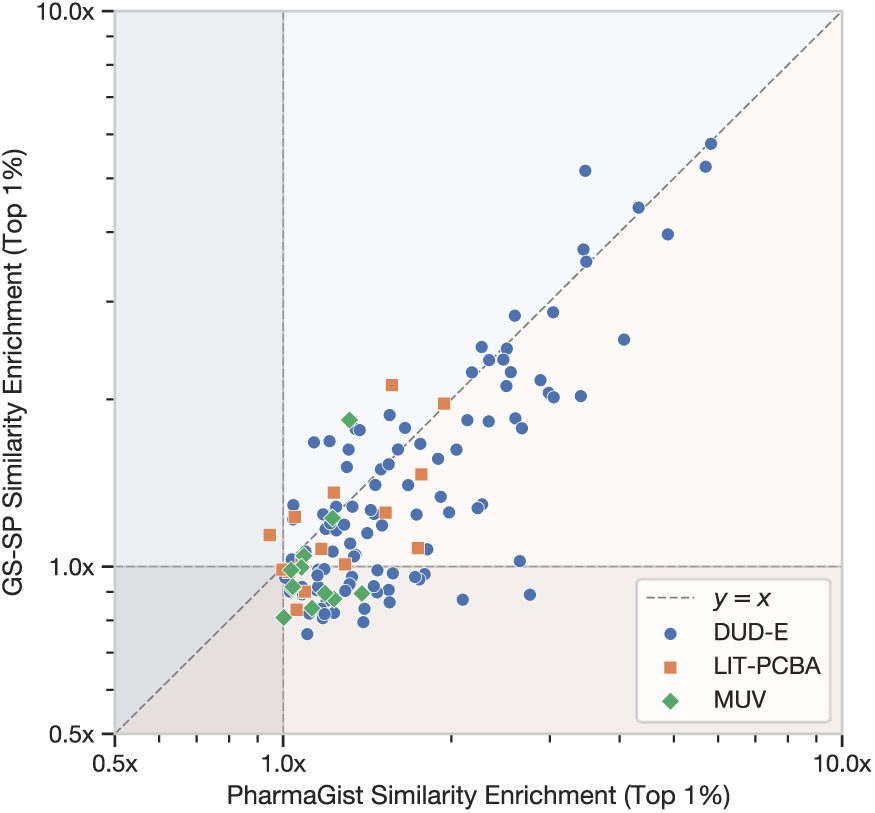
G-screen exhibits lower similarity enrichment in the top 1% ranked molecules compared to PharmaGist. Enrichment for each target is defined as the ratio of the average ECFP4 Tanimoto similarity of the top 1% ranked molecules to that of all actives and decoys of the target.

To illustrate the mechanism behind this capacity to discover less similar molecules, we examined a representative case from the MUV-548 dataset (**Figure 9**). Although its specific rank (top 5.4%) falls outside the strictest early-enrichment cutoffs, this example highlights the qualitative differences in how the two methods score scaffold-divergent compounds. The active molecule (PubChem CID: 2422341) was prioritized much higher by G-screen (top 5.4%) than by PharmaGist (top 57.0%), despite sharing very little structural similarity with the crystal ligand (ECFP4 Tanimoto similarity 0.05). G-screen recognized that the aligned pose preserves key hydrogen-bonding interactions within the protein pocket. In contrast, ligand-based pharmacophore detection (PharmaGist) identified multiple ligand-derived features without the context to distinguish which interactions were structurally essential. This case demonstrates how incorporating protein interaction geometry enables the recovery of active, scaffold- hopped molecules that would be heavily penalized by ligand similarity alone.

**Figure 9.**
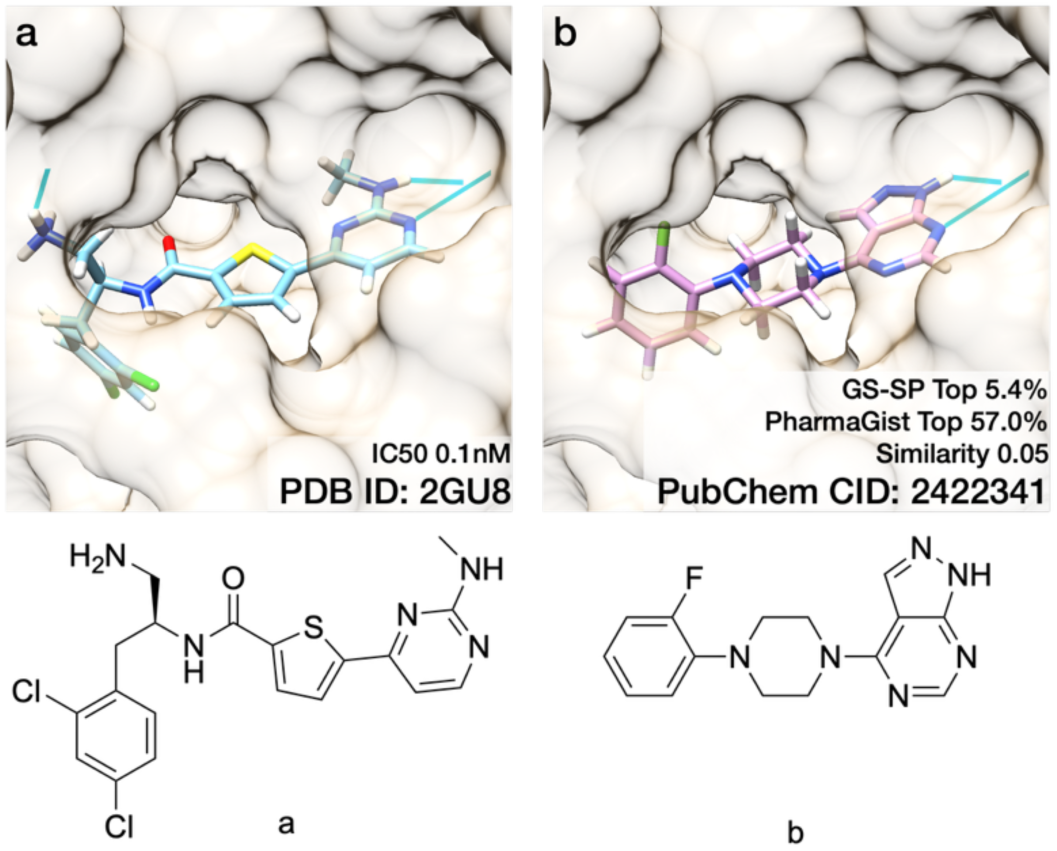
Illustration of protein-aware scoring on the MUV-548 target. The active molecule (b) was ranked within the top 5.4% by G-screen, compared to the top 57.0% by PharmaGist. (a) Crystal ligand and (b) representative active molecule. Hydrogen bonds are shown as sky-blue pseudo-bonds. Residue 125-138 are omitted for clarity.

Additional examples from the DUD-E dataset further clarify the interaction-level behavior of G- screen (**Figure 10**). For the XIAP target (PDB ID: 3HL5), GS-SP achieved an AUROC of 0.97 compared to 0.72 for PharmaGist. The representative active molecule (ChEMBL ID: CHEMBL458742) and decoy (ZINC ID: ZINC4337352) revealed that G-screen correctly prioritized the active compound by preserving hydrogen-bond interactions consistent with the reference complex, while deprioritizing a decoy lacking compatible interaction geometry (**Figure 10**a-b). In this case, protein-aware pharmacophore matching emphasizes interaction hot spots rather than simply counting ligand-derived features.

**Figure 10.**
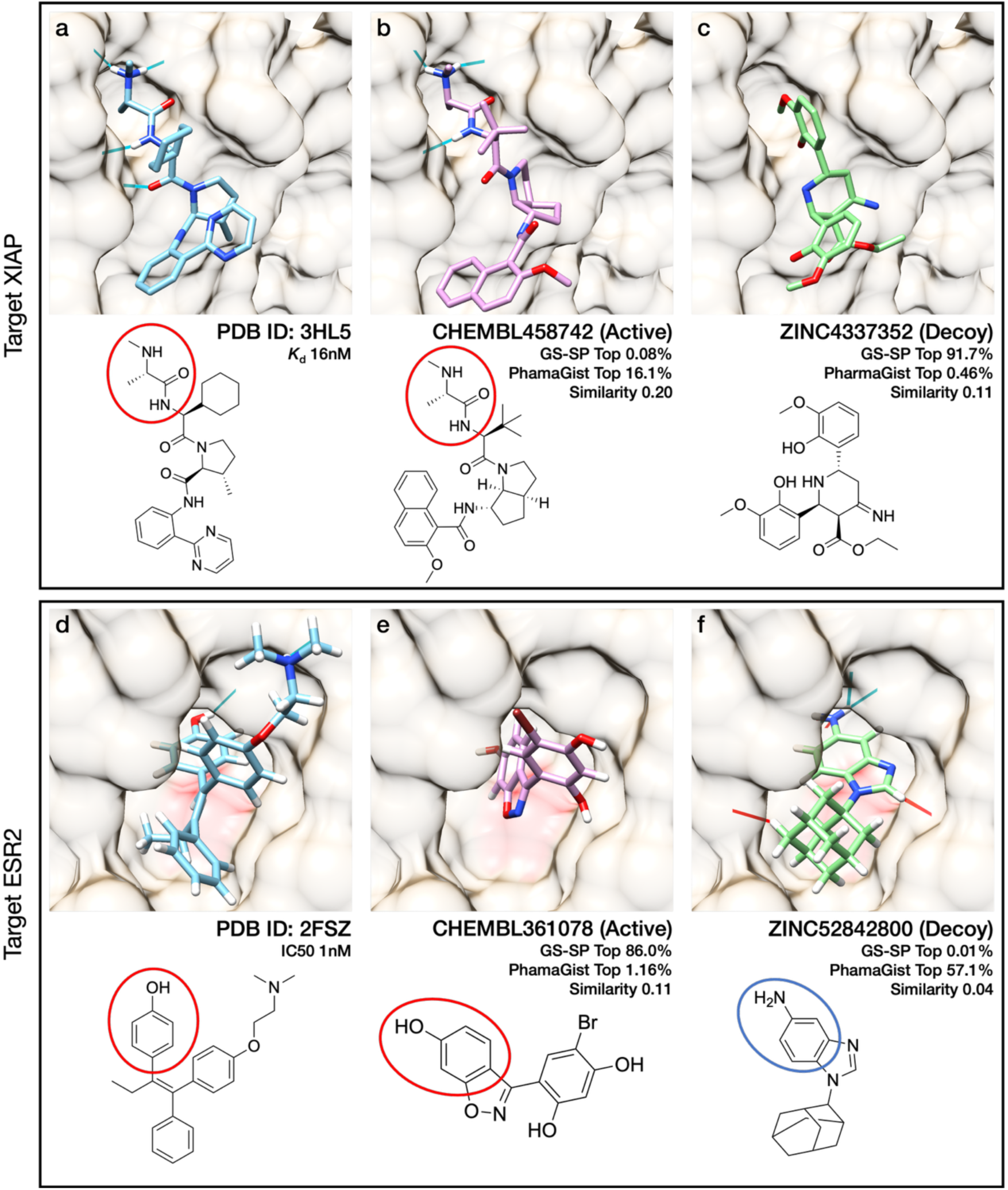
Crystal structures and representative active and decoy molecules for the DUD-E targets XIAP (a–c) and ESR2 (d–f). For each target, the active and decoy molecules were aligned to the corresponding crystal protein– ligand complex using G-align. Hydrogen bonds are shown as sky-blue pseudo-bonds; detected clashes are shown as red pseudo-bonds. A key aromatic moiety of the reference compound and the corresponding moiety of the active compound that participates in a π–π interaction within the reference complex are indicated by red circles. (d–e). Nonessential hydrogen atoms and selected residues (191–209 in d–f) are omitted for clarity.

The ESR2 target (PDB ID: 2FSZ) provides a contrasting example that highlights a limitation of alignment-dependent screening when evaluating protein-ligand interactions. For this target, G-screen achieved an AUROC of 0.83, whereas PharmaGist reached 0.94. The representative active (ChEMBL ID: CHEMBL361078) and decoy (ZINC ID: ZINC52842800) illustrate that G-align did not correctly position a key aromatic moiety of the active compound to reproduce the π–π interaction observed in the reference complex (**Figure 10**d-f). Consequently, protein-aware scoring penalized the active molecule, while a decoy that aligned more closely in shape received a higher score. This example underscores that accurate structural alignment is a prerequisite for effective protein-aware evaluation.

#### 3.3.2. Influence of alignment quality and ligand flexibility on G-screen performance

Because G-screen is an alignment-guided method, its performance depends on whether active molecules align well relative to the screened set, as illustrated by ESR2 (**Figure 10**d-f). To assess this behavior systematically, we defined relative shape similarity of actives as the mean GS-S score of active molecules divided by the mean GS-S score of all screened molecules for each target. The mean GS-S score across the screened set provides a target-specific baseline for alignment to the reference ligand, allowing us to measure whether active molecules align better than expected for that target. Across the main benchmark sets, actives that are more similar to the reference ligand tend to show higher relative shape similarity of actives (**Figure 11**a), and target-level screening performance is higher when relative shape similarity of actives is greater (**Figure 11**b). These results suggest that the G-screen performance depends on the quality of the generated pose.

**Figure 11.**
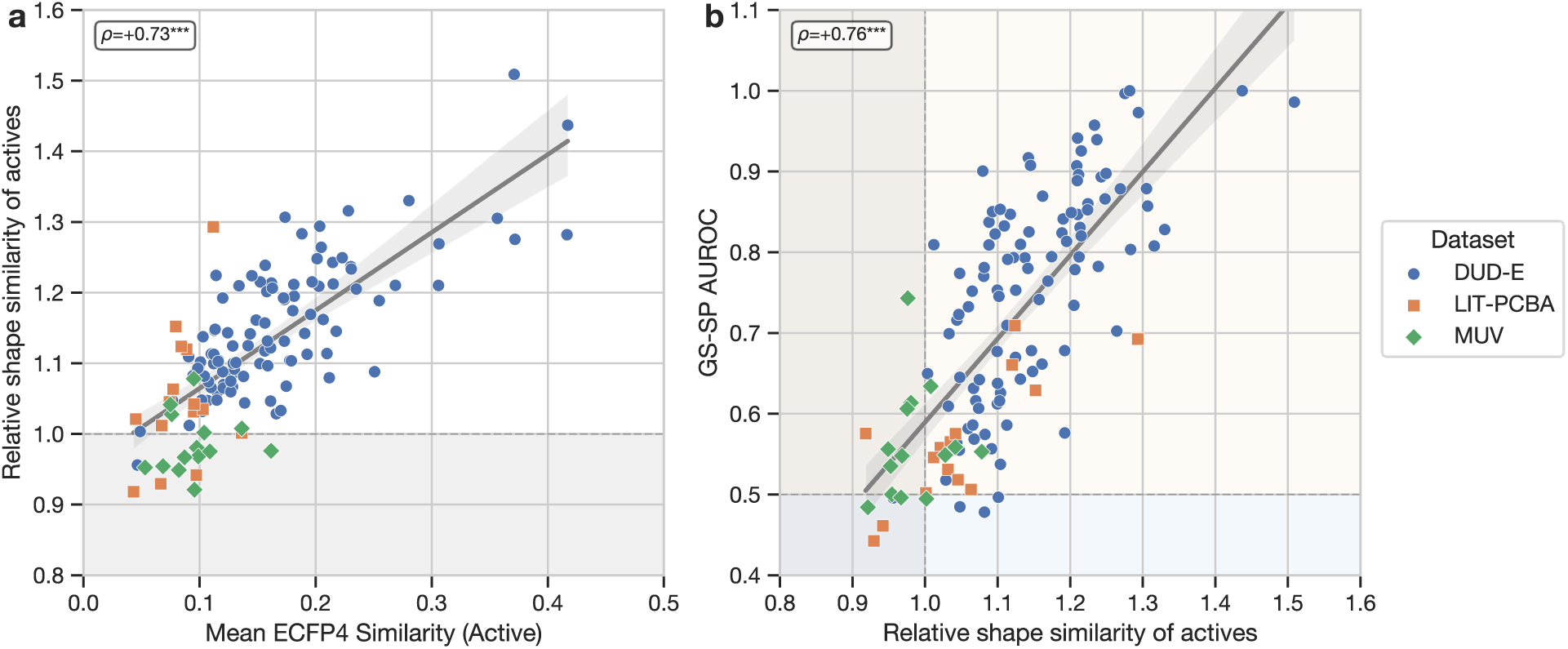
Relationship between alignment quality and G-screen performance. (a) Across targets, greater mean ECFP4 Tanimoto similarity of actives to the reference ligand is associated with higher relative shape similarity of actives. Relative shape similarity of actives is defined as the mean GS-S score of actives divided by the mean GS-S score of all screened molecules per target; values greater than 1 indicate that actives align better than the screened- set average for that target. (b) Across targets, higher relative shape similarity of actives is associated with better screening performance. Each point represents one benchmark target in both panels. Spearman’s *ρ* = 0.73 in (a) and *ρ* = 0.76 in (b); both *p* < 0.001.

To assess the effect of ligand flexibility on G-screen performance, we further analyzed target-level screening performance as a function of ligand flexibility across the benchmark sets. AUROC showed a weak positive association with the raw number of rotatable bonds, whereas this trend became weak or disappeared after normalization by molecular size, suggesting that the apparent rotatable-bond dependence is more likely related to ligand size or complexity than to flexibility itself (**Figure S4**). Consistently, stratified analysis across low (0-5 rotatable bonds), medium (6-7 rotatable bonds), and high (8 or more rotatable bonds) flexibility bins did not reveal a systematic reduction in AUROC or EF1% for GS-SP or the other G-screen variants (**Figure S5**). These results indicate that, within the benchmarked range, G-screen does not exhibit a marked performance penalty for more flexible ligands.

We also examined the sensitivity of the G-screen framework to the alignment method by rescoring poses generated by PharmaGist, whose alignment procedure considers pharmacophore features in addition to molecular shape. A subset of 13 targets spanning the three benchmark datasets and covering a range of GS-P performance levels was selected for this analysis (**Method S5**). In this benchmark, AUROC was similar between G-align-aligned and PharmaGist-aligned poses (0.686 versus 0.668), whereas EF1% was noticeably higher for G-align-aligned poses (15.28 versus 9.29). These results indicate that overall ranking performance is relatively robust to the alignment method, whereas early enrichment depends more strongly on the quality of the alignment procedure (**Figure S6**).

## 4. Conclusions

We have developed G-screen, a scalable protein-aware virtual screening framework that bridges the gap between ligand-based alignment methods and docking-based structure-based screening. By combining fast and flexible ligand alignment (G-align) with protein-aware pharmacophore interaction scoring at atomic resolution, G-screen achieves a favorable balance between computational efficiency, interaction-level interpretability, and practical scalability for ultra-large virtual screening applications.

Across the DUD-E, LIT-PCBA, and MUV benchmarks, G-screen demonstrated competitive discrimination performance while maintaining millisecond-scale per-molecule runtimes under multithreaded execution. Interaction-level case analyses further clarified that protein-aware scoring enables recovery of structurally diverse active molecules beyond simple ligand similarity, while remaining dependent on alignment accuracy and the informativeness of the reference complex. Additional analyses demonstrated that G-screen remains practically applicable within hierarchical large-scale screening workflows and can retain competitive performance even when predicted reference structures are used.

Although G-screen does not replace full docking approaches in modeling induced-fit effects or complex binding energetics, it provides a practical and computationally scalable filtering step prior to more computationally intensive refinement methods when a reference complex structure is available. In this regime, G-screen occupies an intermediate niche between conventional ligand-based virtual screening and computationally intensive docking workflows by combining the scalability and robustness of ligand-based approaches with explicit protein-aware interaction analysis.

We anticipate that G-screen will facilitate scalable structure-guided ligand discovery in both academic and industrial settings. To support broad adoption and reproducibility, G-screen and G-align are freely available at https://github.com/seoklab/gscreen and https://github.com/seoklab/galign, respectively.

## 5. List of Abbreviations

VS: Virtual screening
LBVS: Ligand-based virtual screening
SBVS: Structure-based virtual screening
DOF: Degree of freedom
DUD-E: Directories of Useful Decoys, Enhanced
MUV: Maximum Unbiased Validation
RMSD: Root mean square deviation
PDB: Protein Data Bank
AUROC: Area under the receiver operating characteristic curve
EF: Enrichment factor
AF3: AlphaFold 3

## 6. Declarations

### 6.1. Availability of data and materials

The G-align and G-screen source code used in this article are available at https://github.com/seoklab/galign and https://github.com/seoklab/gscreen, respectively. The datasets supporting the conclusions of this article are included within the article (and its additional files).

### 6.4. Authors’ contributions

J.Y. and C.S. conceived and design the overall study. N.J. developed the algorithms and implemented the code for G-align and G-screen, and conducted all benchmark evaluations. C.S. and H.P. provided critical scientific insights and guidance throughout the research. N.J., C.S., and J.Y. analyzed the results and drafted the manuscript. All authors read and approved the final manuscript.

## References

(1) Bajorath, J. Integration of virtual and high-throughput screening. Nature Reviews Drug Discovery 2002, 1 (11), 882–894. DOI: 10.1038/nrd941.

(2) C Braga, R.; M Alves, V.; C Silva, A.; N Nascimento, M.; C Silva, F.; M Liao, L.; H Andrade, C. Virtual screening strategies in medicinal chemistry: the state of the art and current challenges. Current topics in medicinal chemistry 2014, 14 (16), 1899–1912.

(3) Sabe, V. T.; Ntombela, T.; Jhamba, L. A.; Maguire, G. E. M.; Govender, T.; Naicker, T.; Kruger, H. G. Current trends in computer aided drug design and a highlight of drugs discovered via computational techniques: A review. European Journal of Medicinal Chemistry 2021, 224, 113705. DOI: 10.1016/j.ejmech.2021.113705.

(4) Willett, P. Similarity-based virtual screening using 2D fingerprints. Drug Discovery Today 2006, 11 (23), 1046–1053. DOI: 10.1016/j.drudis.2006.10.005.

(5) Wolber, G.; Langer, T. LigandScout: 3-D Pharmacophores Derived from Protein-Bound Ligands and Their Use as Virtual Screening Filters. Journal of Chemical Information and Modeling 2005, 45 (1), 160–169. DOI: 10.1021/ci049885e.

(6) Kitchen, D. B.; Decornez, H.; Furr, J. R.; Bajorath, J. Docking and scoring in virtual screening for drug discovery: methods and applications. Nature Reviews Drug Discovery 2004, 3 (11), 935–949. DOI: 10.1038/nrd1549.

(7) Shoichet, B. K.; Kobilka, B. K. Structure-based drug screening for G-protein-coupled receptors. Trends in Pharmacological Sciences 2012, 33 (5), 268–272. DOI: 10.1016/j.tips.2012.03.007.

(8) Congreve, M.; Graaf, C. d.; Swain, N. A.; Tate, C. G. Impact of GPCR Structures on Drug Discovery. Cell 2020, 181 (1), 81–91. DOI: 10.1016/j.cell.2020.03.003.

(9) Sadybekov, A. A.; Sadybekov, A. V.; Liu, Y.; Iliopoulos-Tsoutsouvas, C.; Huang, X.-P.; Pickett, J.; Houser, B.; Patel, N.; Tran, N. K.; Tong, F.;, et al. Synthon-based ligand discovery in virtual libraries of over 11 billion compounds. Nature 2022, 601 (7893), 452–459. DOI: 10.1038/s41586-021-04220-9.

(10) Sousa, S. F.; Fernandes, P. A.; Ramos, M. J. Protein–ligand docking: Current status and future challenges. Proteins: Structure, Function, and Bioinformatics 2006, 65 (1), 15–26. DOI: 10.1002/prot.21082.

(11) Buonfiglio, R.; Recanatini, M.; Masetti, M. Protein Flexibility in Drug Discovery: From Theory to Computation. ChemMedChem 2015, 10 (7), 1141–1148. DOI: 10.1002/cmdc.201500086.

(12) Kwon, S.; Seok, C. CSAlign and CSAlign-Dock: Structure alignment of ligands considering full flexibility and application to protein–ligand docking. Computational and Structural Biotechnology Journal 2023, 21, 1–10. DOI: 10.1016/j.csbj.2022.11.047.

(13) Gorgulla, C.; Boeszoermenyi, A.; Wang, Z.-F.; Fischer, P. D.; Coote, P. W.; Padmanabha Das, K. M.; Malets, Y. S.; Radchenko, D. S.; Moroz, Y. S.; Scott, D. A.;, et al. An open-source drug discovery platform enables ultra-large virtual screens. Nature 2020, 580 (7805), 663–668. DOI: 10.1038/s41586-020-2117-z.

(14) Graff, D. E.; Shakhnovich, E. I.; Coley, C. W. Accelerating high-throughput virtual screening through molecular pool-based active learning. Chem. Sci. 2021, 12 (22), 7866–7881. DOI: 10.1039/D0SC06805E.

(15) Zhou, G.; Rusnac, D.-V.; Park, H.; Canzani, D.; Nguyen, H. M.; Stewart, L.; Bush, M. F.; Nguyen, P. T.; Wulff, H.; Yarov-Yarovoy, V.;, et al. An artificial intelligence accelerated virtual screening platform for drug discovery. Nature Communications 2024, 15 (1), 7761. DOI: 10.1038/s41467-024-52061-7.

(16) Corso, G.; Stärk, H.; Jing, B.; Barzilay, R.; Jaakkola, T. DiffDock: Diffusion Steps, Twists, and Turns for Molecular Docking. 2022; p arXiv:2210.01776.

(17) Lu, W.; Zhang, J.; Huang, W.; Zhang, Z.; Jia, X.; Wang, Z.; Shi, L.; Li, C.; Wolynes, P. G.; Zheng, S. DynamicBind: predicting ligand-specific protein-ligand complex structure with a deep equivariant generative model. Nature Communications 2024, 15 (1), 1071. DOI: 10.1038/s41467-024-45461-2.

(18) Cao, D.; Chen, M.; Zhang, R.; Wang, Z.; Huang, M.; Yu, J.; Jiang, X.; Fan, Z.; Zhang, W.; Zhou, H.;, et al. SurfDock is a surface-informed diffusion generative model for reliable and accurate protein–ligand complex prediction. Nature Methods 2025, 22 (2), 310–322. DOI: 10.1038/s41592-024-02516-y.

(19) McNutt, A. T.; Francoeur, P.; Aggarwal, R.; Masuda, T.; Meli, R.; Ragoza, M.; Sunseri, J.; Koes, D. R. GNINA 1.0: molecular docking with deep learning. Journal of Cheminformatics 2021, 13 (1), 43. DOI: 10.1186/s13321-021-00522-2.

(20) Abramson, J.; Adler, J.; Dunger, J.; Evans, R.; Green, T.; Pritzel, A.; Ronneberger, O.; Willmore, L.; Ballard, A. J.; Bambrick, J.;, et al. Accurate structure prediction of biomolecular interactions with AlphaFold 3. Nature 2024, 630 (8016), 493–500. DOI: 10.1038/s41586-024-07487-w.

(21) Wohlwend, J.; Corso, G.; Passaro, S.; Getz, N.; Reveiz, M.; Leidal, K.; Swiderski, W.; Atkinson, L.; Portnoi, T.; Chinn, I.; et al. Boltz-1 Democratizing Biomolecular Interaction Modeling. bioRxiv 2025, 2024.2011.2019.624167. DOI: 10.1101/2024.11.19.624167.

(22) Passaro, S.; Corso, G.; Wohlwend, J.; Reveiz, M.; Thaler, S.; Somnath, V. R.; Getz, N.; Portnoi, T.; Roy, J.; Stark, H.;, et al. Boltz-2: Towards Accurate and Efficient Binding Affinity Prediction. bioRxiv 2025, 2025.2006.2014.659707. DOI: 10.1101/2025.06.14.659707.

(23) Discovery, C.; Boitreaud, J.; Dent, J.; McPartlon, M.; Meier, J.; Reis, V.; Rogozhnikov, A.; Wu, K. Chai-1: Decoding the molecular interactions of life. bioRxiv 2024, 2024.2010.2010.615955. DOI: 10.1101/2024.10.10.615955.

(24) Jia, Y.; Gao, B.; Tan, J.; Zheng, J.; Hong, X.; Zhu, W.; Tan, H.; Xiao, Y.; Tan, L.; Cai, H.;, et al. Deep contrastive learning enables genome-wide virtual screening. Science 2026, 391 (6781), eads9530. DOI: doi:10.1126/science.ads9530.

(25) Gentile, F.; Agrawal, V.; Hsing, M.; Ton, A.-T.; Ban, F.; Norinder, U.; Gleave, M. E.; Cherkasov, A. Deep Docking: A Deep Learning Platform for Augmentation of Structure Based Drug Discovery. ACS Central Science 2020, 6 (6), 939–949. DOI: 10.1021/acscentsci.0c00229.

(26) Mysinger, M. M.; Carchia, M.; Irwin, J. J.; Shoichet, B. K. Directory of Useful Decoys, Enhanced (DUD-E): Better Ligands and Decoys for Better Benchmarking. Journal of Medicinal Chemistry 2012, 55 (14), 6582–6594. DOI: 10.1021/jm300687e.

(27) Tran-Nguyen, V.-K.; Jacquemard, C.; Rognan, D. LIT-PCBA: An Unbiased Data Set for Machine Learning and Virtual Screening. Journal of Chemical Information and Modeling 2020, 60 (9), 4263–4273. DOI: 10.1021/acs.jcim.0c00155.

(28) Rohrer, S. G.; Baumann, K. Maximum Unbiased Validation (MUV) Data Sets for Virtual Screening Based on PubChem Bioactivity Data. Journal of Chemical Information and Modeling 2009, 49 (2), 169–184. DOI: 10.1021/ci8002649.

(29) Schneidman-Duhovny, D.; Dror, O.; Inbar, Y.; Nussinov, R.; Wolfson, H. J. PharmaGist: a webserver for ligand-based pharmacophore detection. Nucleic Acids Research 2008, 36 (suppl_2), W223–W228. DOI: 10.1093/nar/gkn187.

(30) Trott, O.; Olson, A. J. AutoDock Vina: Improving the speed and accuracy of docking with a new scoring function, efficient optimization, and multithreading. Journal of Computational Chemistry 2010, 31 (2), 455–461. DOI: 10.1002/jcc.21334.

(31) Eberhardt, J.; Santos-Martins, D.; Tillack, A. F.; Forli, S. AutoDock Vina 1.2.0: New Docking Methods, Expanded Force Field, and Python Bindings. Journal of Chemical Information and Modeling 2021, 61 (8), 3891–3898. DOI: 10.1021/acs.jcim.1c00203.

(32) Kim, H.; Ryu, S.; Jung, N.; Yang, J.; Seok, C. CSearch: chemical space search via virtual synthesis and global optimization. Journal of Cheminformatics 2024, 16 (1), 137. DOI: 10.1186/s13321-024-00936-8.

(33) Yang, J.; Baek, M.; Seok, C. GalaxyDock3: Protein–ligand docking that considers the full ligand conformational flexibility. Journal of Computational Chemistry 2019, 40 (31), 2739–2748. DOI: 10.1002/jcc.26050.

(34) Nelder, J. A.; Mead, R. A Simplex Method for Function Minimization. The Computer Journal 1965, 7 (4), 308–313. DOI: 10.1093/comjnl/7.4.308.

(35) Hu, J.; Liu, Z.; Yu, D.-J.; Zhang, Y. LS-align: an atom-level, flexible ligand structural alignment algorithm for high-throughput virtual screening. Bioinformatics 2018, 34 (13), 2209–2218. DOI: 10.1093/bioinformatics/bty081.

(36) O’Boyle, N. M.; Banck, M.; James, C. A.; Morley, C.; Vandermeersch, T.; Hutchison, G. R. Open Babel: An open chemical toolbox. Journal of Cheminformatics 2011, 3 (1), 33. DOI: 10.1186/1758-2946-3-33.

(37) Free Software Foundation, I. GNU Time. 1996.

(38) Mills, J. E. J.; Dean, P. M. Three-dimensional hydrogen-bond geometry and probability information from a crystal survey. Journal of Computer-Aided Molecular Design 1996, 10 (6), 607–622. DOI: 10.1007/BF00134183.

(39) Pettersen, E. F.; Goddard, T. D.; Huang, C. C.; Couch, G. S.; Greenblatt, D. M.; Meng, E. C.; Ferrin, T. E. UCSF Chimera—A visualization system for exploratory research and analysis. Journal of Computational Chemistry 2004, 25 (13), 1605–1612. DOI: 10.1002/jcc.20084.

(40) Greene, J.; Kahn, S.; Savoj, H.; Sprague, P.; Teig, S. Chemical Function Queries for 3D Database Search. Journal of Chemical Information and Computer Sciences 1994, 34 (6), 1297–1308. DOI: 10.1021/ci00022a012.

(41) McGaughey, G. B.; Gagné, M.; Rappé, A. K. π-Stacking Interactions: ALIVE AND WELL IN PROTEINS*. Journal of Biological Chemistry 1998, 273 (25), 15458–15463. DOI: 10.1074/jbc.273.25.15458.

(42) Da Silva, F.; Desaphy, J.; Rognan, D. IChem: A Versatile Toolkit for Detecting, Comparing, and Predicting Protein–Ligand Interactions. ChemMedChem 2018, 13 (6), 507–510. DOI: 10.1002/cmdc.201700505 (accessed 2026/04/15).

(43) Tran-Nguyen, V.-K.; Da Silva, F.; Bret, G.; Rognan, D. All in One: Cavity Detection, Druggability Estimate, Cavity-Based Pharmacophore Perception, and Virtual Screening. Journal of Chemical Information and Modeling 2019, 59 (1), 573–585. DOI: 10.1021/acs.jcim.8b00684.

(44) Waskom, M. L. seaborn: statistical data visualization. Journal of Open Source Software 2021, 6 (60), 3021. DOI: 10.21105/joss.03021.

(45) Hunter, J. D. Matplotlib: A 2D graphics environment. Computing in Science & Engineering 2007, 9 (3), 90–95. DOI: 10.1109/MCSE.2007.55.

(46) Harris, C. R.; Millman, K. J.; Walt, S. J. v. d.; Gommers, R.; Virtanen, P.; Cournapeau, D.; Wieser, E.; Taylor, J.; Berg, S.; Smith, N. J.;, et al. Array programming with NumPy. Nature 2020, 585 (7825), 357–362. DOI: 10.1038/s41586-020-2649-2.

(47) Virtanen, P.; Gommers, R.; Oliphant, T. E.; Haberland, M.; Reddy, T.; Cournapeau, D.; Burovski, E.; Peterson, P.; Weckesser, W.; Bright, J.;, et al. SciPy 1.0: Fundamental Algorithms for Scientific Computing in Python. Nature Methods 2020, 17, 261–272. DOI: 10.1038/s41592-019-0686-2.

(48) team, T. p. d. pandas-dev/pandas: Pandas. Zenodo: 2020.

(49) McKinney, W. Data Structures for Statistical Computing in Python. 2010, 2010; Walt, S. v. d., Millman, J., Eds.; pp 56–61. DOI: 10.25080/Majora-92bf1922-00a.

(50) Pedregosa, F.; Varoquaux, G.; Gramfort, A.; Michel, V.; Thirion, B.; Grisel, O.; Blondel, M.; Prettenhofer, P.; Weiss, R.; Dubourg, V.;, et al. Scikit-learn: Machine Learning in Python. Journal of Machine Learning Research 2011, 12, 2825–2830.

(51) Granger, B. E.; Pérez, F. Jupyter: Thinking and Storytelling With Code and Data. Computing in Science & Engineering 2021, 23 (2), 7–14. DOI: 10.1109/MCSE.2021.3059263.

(52) Hagberg, A. A.; Schult, D. A.; Swart, P. J. Exploring Network Structure, Dynamics, and Function using NetworkX. 2008, 2008; Varoquaux, G., Vaught, T., Millman, J., Eds.; Pasadena, CA USA, pp 11–15.

