## Supplementary Information for "G-screen: Scalable Protein-Aware Virtual Screening through Flexible Ligand Alignment"

**For**

**Method S1. Definition of Charged Interaction**

Charged interaction features were assigned following the LigandScout[1] definition. Positively ionizable features were defined as atoms or functional groups expected to be protonated at physiological pH, including basic amines, primary and secondary amidines, guanidines, and formal positive charges not neutralized by adjacent negative charges. Negatively ionizable features were defined as atoms or groups expected to be deprotonated at physiological pH, including sulfonic, phosphonic, sulfinic, carboxylic, and phosphinic acids, trifluoromethyl sulfonamide hydrogens, tetrazoles, and formal negative charges not neutralized by adjacent positive charges. A positive or negative ionizable feature was assigned when a complementary oppositely charged feature was present on the protein side within 1.5 Å to 5.6 Å; the same distance criterion was also used to define a match.

Protein-side charged groups were defined from standard charged residues, namely arginine, histidine, and lysine for positive features, and aspartic acid and glutamic acid for negative features. Ligand-side charged groups were identified by sequential application of the predefined criteria using SMARTS pattern matching in OpenBabel.

**Method S2. Construction and statistical evaluation of the cross-template validation subset from LIT-PCBA**

To enable a more rigorous evaluation of performance differences among the benchmarked methods, we constructed an additional cross-template validation subset from LIT-PCBA, a benchmark that provides multiple experimentally determined protein–ligand complex structures for many targets. Depending on the target, LIT-PCBA contains between 1 and 15 candidate reference complexes derived from the Protein Data Bank (PDB). Direct use of all available reference complexes could overweight highly similar reference ligands when structurally redundant templates are overrepresented for a given target. To reduce such redundancy while preserving structural diversity, we constructed a clustered cross-template subset for each target. This redundancy was non-negligible for some targets; for example, for the *ESR1_ant* target, 8 of the 15 available reference ligands were assigned to a single similarity cluster.

Reference ligands from the full LIT-PCBA dataset were clustered separately for each target using Tanimoto distance computed from ECFP4 fingerprints, and at most five reference complexes were retained per target. Representative reference ligands were selected as cluster medoids, except when a cluster contained the reference complex used in the main benchmark analysis, in which case that complex was retained to maintain consistency with the primary benchmark setting. Active molecules were processed analogously, yielding up to 10 representative actives per target based on cluster medoids. In contrast, the number of decoys in LIT-PCBA is substantially larger, reaching up to approximately 360,000 molecules per target and averaging about 180,000. Therefore, to construct a computationally tractable cross-template validation set, 1,000 decoy molecules were randomly sampled per target rather than clustered.

For cross-template validation, each selected reference complex for a given target was used independently as a template, and screening performance was evaluated using AUROC and EF1%. To compare average performance across methods, metric values were first averaged across the selected reference complexes for each target, yielding one target-level value per method. Planned pairwise comparisons were then performed between each G-screen variant and each baseline method using paired Wilcoxon signed-rank tests across the 13 targets. Corrected $p$ values were obtained using the Benjamini–Hochberg false discovery rate procedure.

To quantify sensitivity to template choice, cross-template dispersion was calculated separately for each target and method. Using the selected reference complexes for each target, dispersion was defined as the standard deviation for AUROC and as the standard deviation of $\log(1 + \text{EF1\%})$ for EF1%. Dispersion values were then compared between G-screen variants and baseline methods using two-sided paired Wilcoxon signed-rank tests across targets, with Benjamini-Hochberg correction applied for multiple comparisons.

**Method S3. Preparation of AF3-predicted reference models**

To assess the applicability of G-screen variants to computationally predicted complex structures in addition to experimentally determined structures, we generated AlphaFold3 (AF3)[2] models for all reference complexes used in the main benchmark analysis.

For each benchmark target with an available reference complex, the Chemical Component Dictionary (CCD) ID of the target ligand and the amino acid sequences of protein chains containing at least one atom within 4.5 Å of the target ligand were identified. Additional small molecules were included if any atom was located within the same 4.5 Å cutoff of the target ligand.

MSA generation was performed with MMseqs2[3] (version 18.8cc5c) using the default ColabFold settings and databases.[4] Template retrieval was performed through the AF3 data pipeline without imposing a template release-date cutoff. Structure prediction with AF3 was conducted using the default inference settings and a single seed. For each target, the top-ranked model according to the AF3 ranking score was selected for downstream analysis. The resulting predicted structure was then separated into a protein structure in standard PDB format and a ligand structure in MOL2 format.

For MUV targets lacking experimental structures at the time of benchmark curation, we additionally generated AF3 models for all active molecules. For each target, the corresponding ligand SMILES strings, together with the full UniProt[5] sequence of the target gene associated with the relevant PubChem[6] assay, were used as AF3 inputs. The inference setup was the same as described above, including MSA generation, template retrieval, and inference parameters. Predicted structures for all active molecules were ranked according to the AF3 ranking score, and the highest-ranked model without detectable intra- or intermolecular clashes, as assessed using UCSF ChimeraX (version 1.10),[7-9] was selected as the reference complex. This selected complex was then prepared in the same manner as described above, yielding protein and ligand files in PDB and MOL2 formats, respectively. The selected active molecule was subsequently removed from the active pool, and benchmarking was performed as described for the other targets. Results for these targets are reported separately for clarity.

**Method S4. Empirical memory-scaling analysis for large-scale virtual screening**

When virtual screening is performed at large scale, practical throughput depends not only on CPU efficiency but also on memory efficiency and memory-scaling behavior, as excessive memory usage can limit vertical scaling to the available CPU resources. To assess the practical feasibility of large-scale parallel execution, we therefore analyzed the memory-scaling behavior of GS-S, GS-P/SP, Flexi-LS-align, and PharmaGist, with particular attention to methods that do not natively support multithreaded or multiprocess execution (Flexi-LS-align and PharmaGist). AutoDock Vina was excluded from this analysis because a best-case estimate assuming perfect linear scaling from the single-thread runtime to 128 threads still yielded per-molecule runtimes at least 100-fold slower than those of GS-SP at 128 threads (**Table 3**), making memory scaling less relevant than raw computational cost for its practical throughput.

Using the same 100-molecule subset employed for the computational efficiency measurements (**Section 2.4.2**), peak memory usage was measured as the maximum resident set size (RSS) reported by GNU Time for runs containing 1, 10, 50, and 100 molecules. For each method, empirical scaling behavior was summarized across targets in terms of both absolute peak memory usage and memory usage normalized to the smallest tested workload (1 query molecule). Based on these measurements, we further estimated the fraction of targets that would remain feasible under a 128-thread execution setting at representative total RAM budgets.

**Method S5. Analysis of alignment-method sensitivity within the G-screen scoring framework**

To assess the dependence of G-screen on the upstream alignment procedure, we performed an additional comparison on a subset of benchmark targets. Because the purpose of this experiment was to isolate the alignment-method sensitivity of the pharmacophore-based scoring component, we used GS-P rather than the proposed GS-SP scoring scheme. GS-SP depends on, and is optimized for, the GS-S alignment score; applying it directly to PharmaGist-generated poses would therefore confound the comparison. Target selection was designed to sample a range of GS-P performance levels while keeping the computational cost of PharmaGist-based alignment tractable.

For the DUD-E dataset, which contains the largest number of targets, targets were first divided into three equally sized bins according to GS-P AUROC. From each bin, the three targets with the smallest combined number of active and decoy molecules were selected, yielding nine targets in total. For the LIT-PCBA and MUV datasets, which contain fewer targets, each dataset was divided into two equally sized bins by GS-P AUROC, and one target with the smallest combined active and decoy set size was selected from each bin. For MUV, two targets on which PharmaGist failed were excluded from candidate selection after binning. This procedure yielded a total of 13 targets spanning DUD-E, LIT-PCBA, and MUV.

Within this subset, the same protein-aware GS-P scoring function was applied to two alternative pose sets: poses generated by the default G-align procedure and poses generated by PharmaGist. Screening performance was then compared in terms of AUROC and EF1%.

**Table S1. Effect of structural and interaction modifications on G-screen performance.**

| Benchmark Dataset | Screening Method | | Screening Performance | | | | | | | |
|---|---|---|---|---|---|---|---|---|---|---|
| | | | AUROC | (Δ) | EF0.1% | (Δ) | EF1% | (Δ) | EF5% | (Δ) |
| DUD-E | GS-P | (Baseline) | 0.72 | | 21.70 | | 13.62 | | 5.86 | |
| | | + Charged Interaction | 0.71 | -0.01 | 20.35 | -1.35 | 13.54 | -0.08 | 5.83 | -0.03 |
| | | AF3 Model Ref. | 0.70 | -0.02 | 15.42 | -6.28 | 10.45 | -3.17 | 4.94 | -0.92 |
| | GS-SP | (Baseline) | 0.76 | | 15.02 | | 14.38 | | 6.92 | |
| | | + Charged Interaction | 0.75 | -0.01 | 14.81 | -0.21 | 14.78 | +0.40 | 7.04 | +0.12 |
| | | AF3 Model Ref. | 0.72 | -0.04 | 11.82 | -3.20 | 9.86 | -4.52 | 5.21 | -1.71 |
| LIT-PCBA | GS-P | (Baseline) | 0.56 | | 6.46 | | 3.78 | | 2.03 | |
| | | + Charged Interaction | 0.57 | +0.01 | 4.07 | -2.39 | 3.15 | -0.63 | 2.08 | +0.05 |
| | | AF3 Model Ref. | 0.56 | ~0 | 12.68 | +6.22 | 3.58 | -0.20 | 1.96 | -0.07 |
| | GS-SP | (Baseline) | 0.56 | | 15.73 | | 4.80 | | 2.20 | |
| | | + Charged Interaction | 0.58 | +0.02 | 7.62 | -8.11 | 4.22 | -0.58 | 2.18 | -0.02 |
| | | AF3 Model Ref. | 0.56 | ~0 | 15.72 | -0.01 | 5.08 | +0.28 | 2.16 | -0.04 |
| MUV | GS-P | (Baseline) | 0.56 | | 1.61 | | 1.94 | | 1.94 | |
| | | + Charged Interaction | 0.56 | ~0 | 0.00 | -1.61 | 1.15 | -0.79 | 1.56 | -0.38 |
| | | AF3 Model Ref. | 0.53 | -0.03 | 1.35 | -0.26 | 1.74 | -0.20 | 1.32 | -0.62 |
| | GS-SP | (Baseline) | 0.56 | | 2.87 | | 2.43 | | 2.21 | |
| | | + Charged Interaction | 0.57 | +0.01 | 0.00 | -2.87 | 1.30 | -1.13 | 1.68 | -0.53 |
| | | AF3 Model Ref. | 0.53 | -0.03 | 1.38 | -1.49 | 2.14 | -0.29 | 1.82 | -0.39 |

**Table S2. Paired Wilcoxon signed-rank test results for the cross-template validation subset of LIT-PCBA.**

(a) Planned pairwise comparisons between G-screen variants with baseline methods across 13 LIT-PCBA targets under the multi-reference protocol (up to 5 clustered references per target). Mean differences are defined as Δ = (G-screen) – (baseline), such that positive values indicate better performance for G-screen.

| Metric | G-screen | Baseline | Mean Performance | | Mean Δ | *p* (corrected)[a] | Significance[b] |
|---|---|---|---|---|---|---|---|
| | | | G-screen | Baseline | | | |
| AUROC | GS-S | Flexi-LS-align | 0.39 | 0.40 | -0.0048 | 0.5879 | |
| | | PharmaGist[c] | | 0.48 | -0.0851 | 0.0308 | * |
| | | AutoDock Vina | | 0.42 | -0.0322 | 0.0904 | |
| | GS-P | Flexi-LS-align | 0.51 | 0.40 | 0.1172 | 0.0073 | ** |
| | | PharmaGist[c] | | 0.48 | 0.0368 | 0.0904 | |
| | | AutoDock Vina | | 0.42 | 0.0898 | 0.0073 | ** |
| | GS-SP | Flexi-LS-align | 0.52 | 0.40 | 0.1185 | 0.0073 | ** |
| | | PharmaGist[c] | | 0.48 | 0.0381 | 0.0597 | |
| | | AutoDock Vina | | 0.42 | 0.0911 | 0.0077 | ** |
| EF1% | GS-S | Flexi-LS-align | 2.12 | 2.82 | -0.7063 | 0.4688 | |
| | | PharmaGist[c] | | 5.72 | -3.6021 | 0.0703 | |
| | | AutoDock Vina | | 2.31 | -0.1883 | 0.8438 | |
| | GS-P | Flexi-LS-align | 4.22 | 2.82 | 1.3916 | 0.2355 | |
| | | PharmaGist[c] | | 5.72 | -1.5042 | 0.5550 | |
| | | AutoDock Vina | | 2.31 | 1.9096 | 0.1230 | |
| | GS-SP | Flexi-LS-align | 4.50 | 2.82 | 1.6747 | 0.1890 | |
| | | PharmaGist[c] | | 5.72 | -1.2211 | 0.6262 | |
| | | AutoDock Vina | | 2.31 | 2.1927 | 0.1890 | |

[a]Corrected *p* values were obtained using the Benjamini-Hochberg false discovery rate procedure.

[b]* $p<0.05$; ** $p<0.01$.

[c]PharmaGist failed on five reference-complex evaluations across three targets because memory usage exceeded 256 GB; those evaluations were excluded from the corresponding comparisons.

(b) Stability analysis based on cross-template dispersion across 13 LIT-PCBA targets. Dispersion was computed for each target and method across clustered reference complexes (up to 5 clustered references per target). Mean differences are reported as Δ = (G-screen) – (baseline), such that positive values indicate greater template sensitivity for G-screen. No difference remained statistically significant after multiple-testing correction, although some comparisons showed numerically higher dispersion for G-screen variants.

| Metric | G-screen | Baseline | Mean Dispersion[a] | | Mean Δ | 95% CI | *p* (corrected)[b] |
|---|---|---|---|---|---|---|---|
| | | | G-screen | Baseline | | | |
| AUROC | GS-S | Flexi-LS-align | 0.0392 | 0.0429 | -0.0037 | [-0.0192, 0.0107] | 0.7501 |
| | | PharmaGist[c] | | 0.0440 | -0.0048 | [-0.0232, 0.0166] | 0.7460 |
| | | AutoDock Vina | | 0.0432 | -0.0040 | [-0.0232, 0.0156] | 0.7460 |
| | GS-P | Flexi-LS-align | 0.0724 | 0.0429 | 0.0295 | [0.0118, 0.0463] | 0.0769 |
| | | PharmaGist[c] | | 0.0440 | 0.0285 | [0.0098, 0.0466] | 0.0773 |
| | | AutoDock Vina | | 0.0432 | 0.0292 | [0.0116, 0.0471] | 0.0769 |
| | GS-SP | Flexi-LS-align | 0.0707 | 0.0429 | 0.0278 | [0.0087, 0.0458] | 0.0769 |
| | | PharmaGist[c] | | 0.0440 | 0.0267 | [0.0072, 0.0470] | 0.0895 |
| | | AutoDock Vina | | 0.0432 | 0.0275 | [0.0097, 0.0461] | 0.0769 |
| EF1% | GS-S | Flexi-LS-align | 0.3731 | 0.7667 | -0.3936 | [-0.6943, -0.0978] | 0.0804 |
| | | PharmaGist[c] | | 0.6534 | -0.2803 | [-0.5992, -0.0408] | 0.0804 |
| | | AutoDock Vina | | 0.3968 | -0.0237 | [-0.3843, 0.3263] | 0.9492 |
| | GS-P | Flexi-LS-align | 0.7397 | 0.7667 | -0.0271 | [-0.4451, 0.3964] | 1.0000 |
| | | PharmaGist[c] | | 0.6534 | 0.0862 | [-0.3923, 0.5367] | 0.7828 |
| | | AutoDock Vina | | 0.3968 | 0.3428 | [0.0234, 0.6707] | 0.2355 |
| | GS-SP | Flexi-LS-align | 0.7669 | 0.7667 | 0.0002 | [-0.4070, 0.4074] | 1.0000 |
| | | PharmaGist[c] | | 0.6534 | 0.1135 | [-0.3639, 0.5543] | 0.7828 |
| | | AutoDock Vina | | 0.3968 | 0.3701 | [0.0584, 0.6938] | 0.1289 |

[a]Mean dispersion was calculated as the standard deviation across reference complexes for AUROC, and the standard deviation of $\log(1 + \mathrm{EF1\%})$ for EF1%.

[b]Corrected $p$ values were obtained using the Benjamini-Hochberg false discovery rate procedure.

[c]PharmaGist failed to run on five reference complexes across three targets due to excessive memory usage exceeding 256GB; those reference-complex evaluations were excluded from the corresponding comparisons.

**Table S3. Virtual screening performance on three MUV targets lacking experimental structures.**

| Screening Method | Screening Performance (↑) | | | | Similarity Enrichment[a] (Top 1%) |
|---|---|---|---|---|---|
| | AUROC | EF0.1% | EF1% | EF5% | |
| GS-S | 0.46 | 0.00 | 0.00 | 0.23 | +28.1% |
| GS-P | <u>0.53</u> | **1.34** | <u>2.55</u> | 1.11 | +9.8% |
| GS-SP | 0.52 | <u>1.16</u> | 2.29 | 0.97 | -1.9% |
| Flexi-LS-align | 0.51 | 0.00 | 1.14 | **1.56** | +40.8% |
| PharmaGist | **0.57** | 0.00 | 1.14 | 0.98 | +26.8% |
| AutoDock Vina | 0.52 | 0.00 | **3.46** | <u>1.33</u> | -0.2% |

[a]Similarity enrichment was defined as the ratio of the average ECFP4 Tanimoto similarity of the top 1% ranked molecules to that of all actives and decoys for a given target.

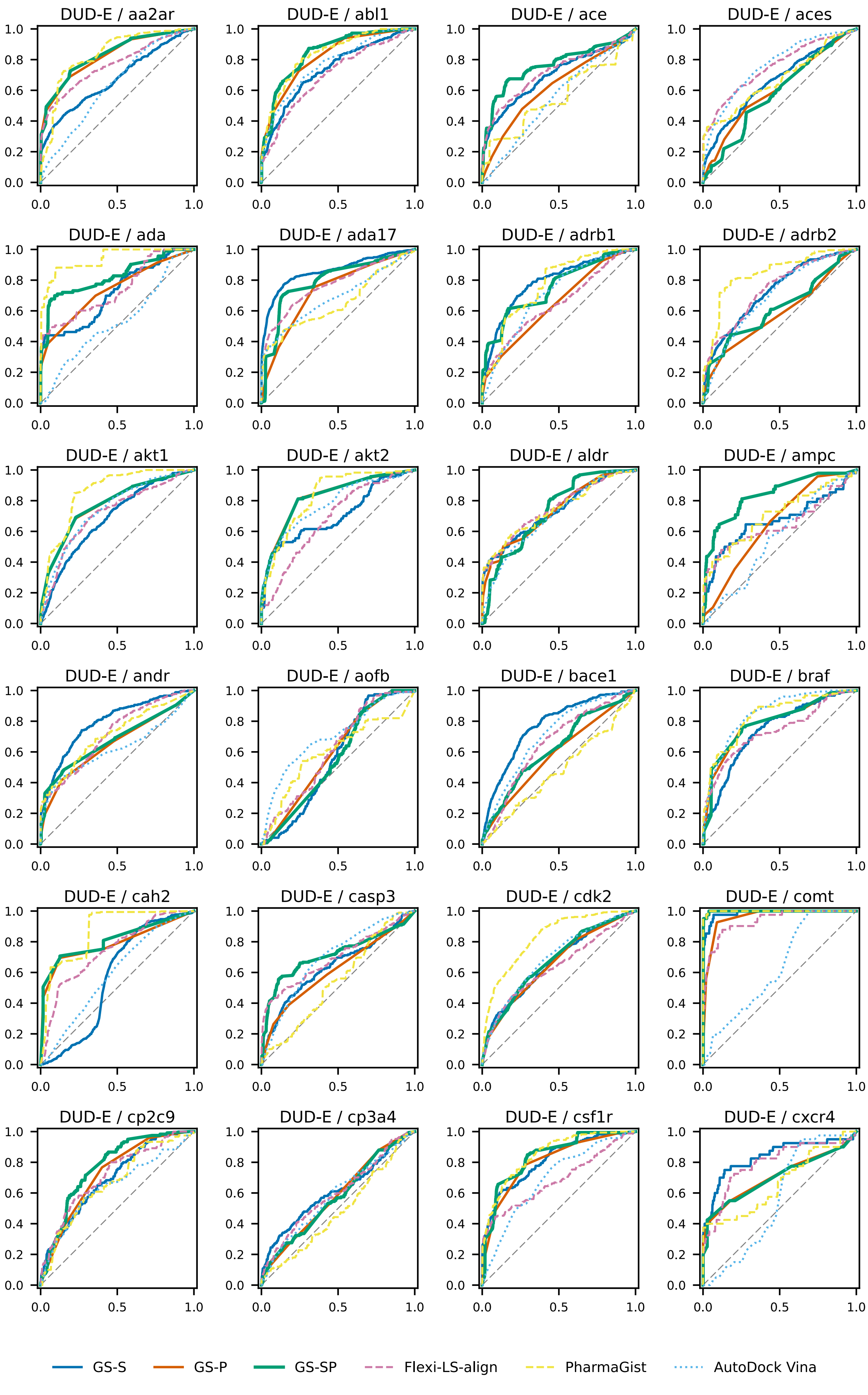
DUD-E / aa2ar
DUD-E / abl1
DUD-E / ace
DUD-E / aces
DUD-E / ada
DUD-E / ada17
DUD-E / adrb1
DUD-E / adrb2
DUD-E / akt1
DUD-E / akt2
DUD-E / aldr
DUD-E / ampc
DUD-E / andr
DUD-E / aofb
DUD-E / bace1
DUD-E / braf
DUD-E / cah2
DUD-E / casp3
DUD-E / cdk2
DUD-E / comt
DUD-E / cp2c9
DUD-E / cp3a4
DUD-E / csf1r
DUD-E / cxcr4
GS-S
GS-P
GS-SP
Flexi-LS-align
PharmaGist
AutoDock Vina

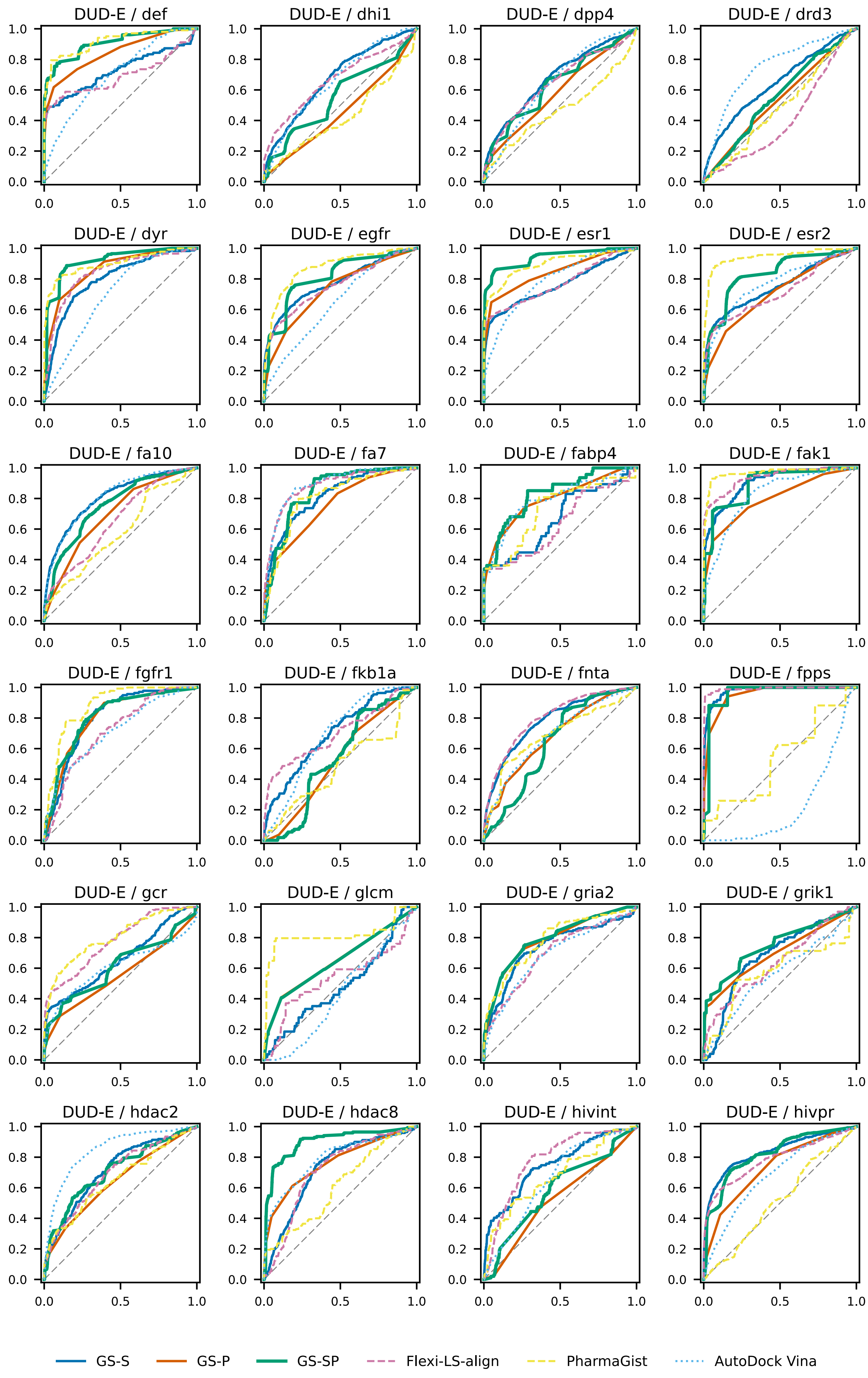
DUD-E / def
DUD-E / dhi1
DUD-E / dpp4
DUD-E / drd3
DUD-E / dyr
DUD-E / egfr
DUD-E / esr1
DUD-E / esr2
DUD-E / fa10
DUD-E / fa7
DUD-E / fabp4
DUD-E / fak1
DUD-E / fgfr1
DUD-E / fkb1a
DUD-E / fnta
DUD-E / fpps
DUD-E / gcr
DUD-E / glcm
DUD-E / gria2
DUD-E / grik1
DUD-E / hdac2
DUD-E / hdac8
DUD-E / hivint
DUD-E / hivpr
GS-S
GS-P
GS-SP
Flexi-LS-align
PharmaGist
AutoDock Vina

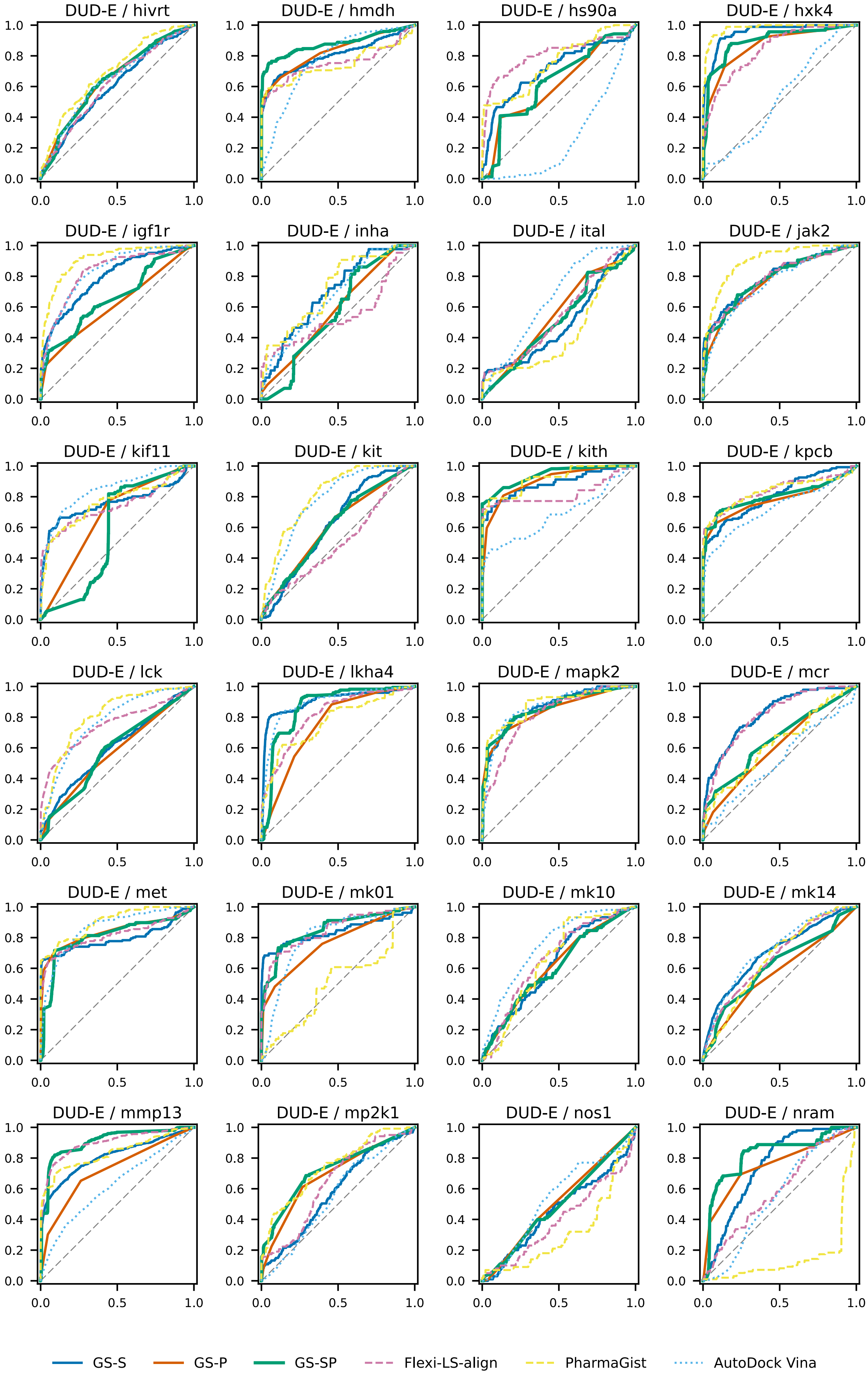
DUD-E / hivrt
DUD-E / hmdh
DUD-E / hs90a
DUD-E / hxk4
DUD-E / igf1r
DUD-E / inha
DUD-E / ital
DUD-E / jak2
DUD-E / kif11
DUD-E / kit
DUD-E / kith
DUD-E / kpcb
DUD-E / lck
DUD-E / lkha4
DUD-E / mapk2
DUD-E / mcr
DUD-E / met
DUD-E / mk01
DUD-E / mk10
DUD-E / mk14
DUD-E / mmp13
DUD-E / mp2k1
DUD-E / nos1
DUD-E / nram
GS-S
GS-P
GS-SP
Flexi-LS-align
PharmaGist
AutoDock Vina

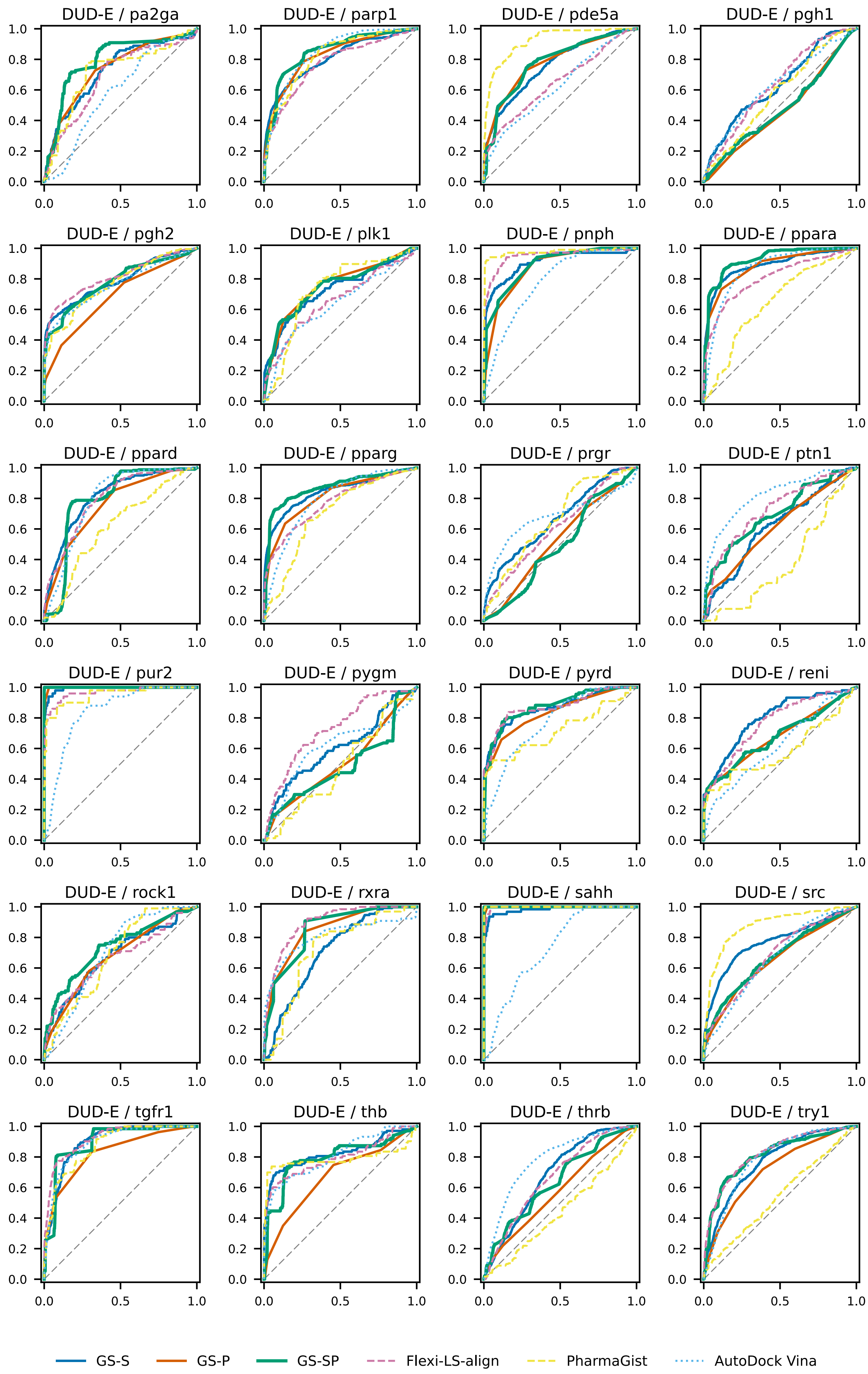
DUD-E / pa2ga
DUD-E / parp1
DUD-E / pde5a
DUD-E / pgh1
DUD-E / pgh2
DUD-E / plk1
DUD-E / pnph
DUD-E / ppara
DUD-E / ppard
DUD-E / pparg
DUD-E / prgr
DUD-E / ptn1
DUD-E / pur2
DUD-E / pygm
DUD-E / pyrd
DUD-E / reni
DUD-E / rock1
DUD-E / rxra
DUD-E / sahh
DUD-E / src
DUD-E / tgfr1
DUD-E / thb
DUD-E / thrb
DUD-E / try1
GS-S
GS-P
GS-SP
Flexi-LS-align
PharmaGist
AutoDock Vina

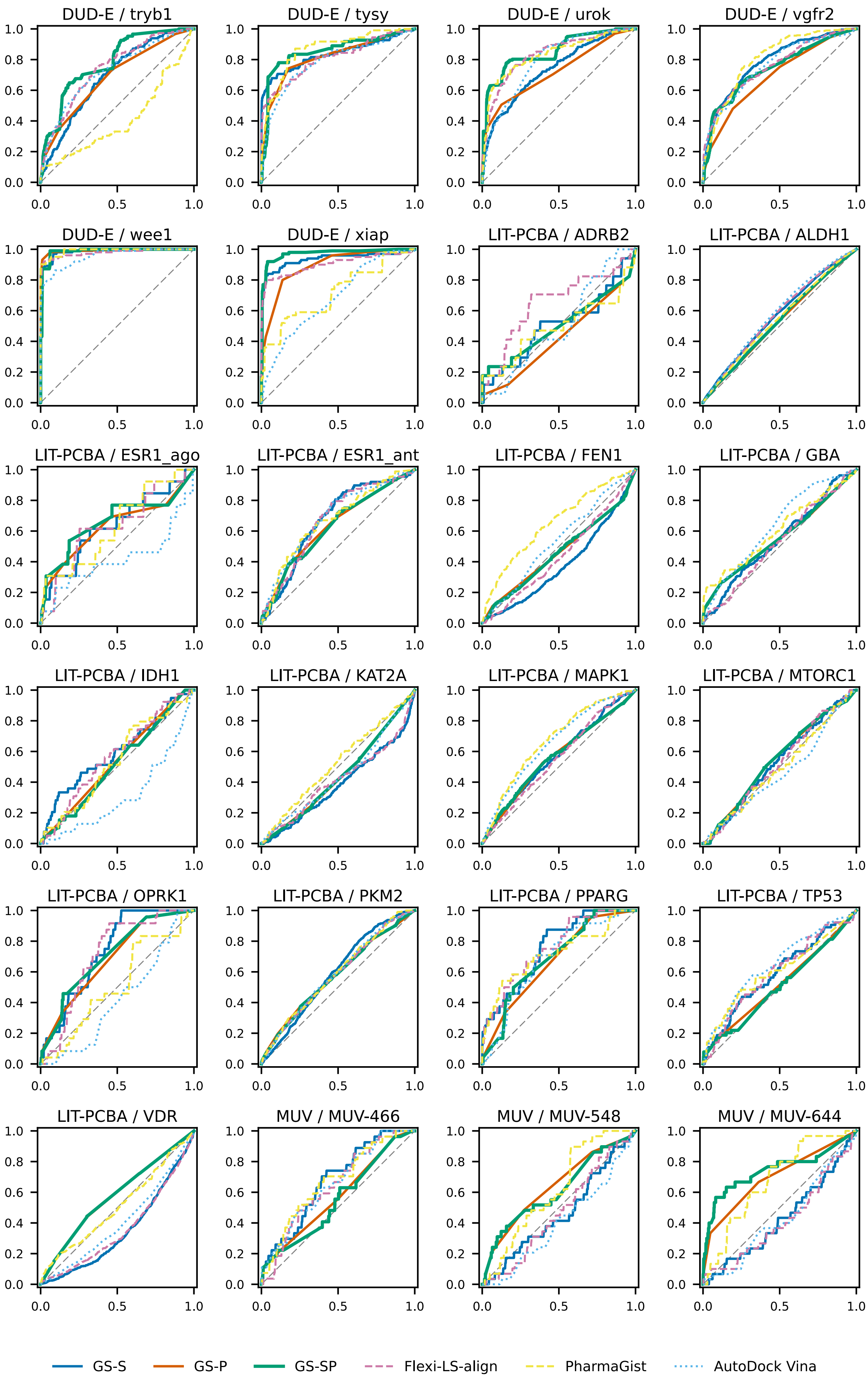

DUD-E / tryb1
DUD-E / tysy
DUD-E / urok
DUD-E / vgfr2
DUD-E / wee1
DUD-E / xiap
LIT-PCBA / ADRB2
LIT-PCBA / ALDH1
LIT-PCBA / ESR1_ago
LIT-PCBA / ESR1_ant
LIT-PCBA / FEN1
LIT-PCBA / GBA
LIT-PCBA / IDH1
LIT-PCBA / KAT2A
LIT-PCBA / MAPK1
LIT-PCBA / MTORC1
LIT-PCBA / OPRK1
LIT-PCBA / PKM2
LIT-PCBA / PPARG
LIT-PCBA / TP53
LIT-PCBA / VDR
MUV / MUV-466
MUV / MUV-548
MUV / MUV-644
0.0
0.2
0.4
0.6
0.8
1.0
0.0
0.5
1.0
GS-S
GS-P
GS-SP
Flexi-LS-align
PharmaGist
AutoDock Vina

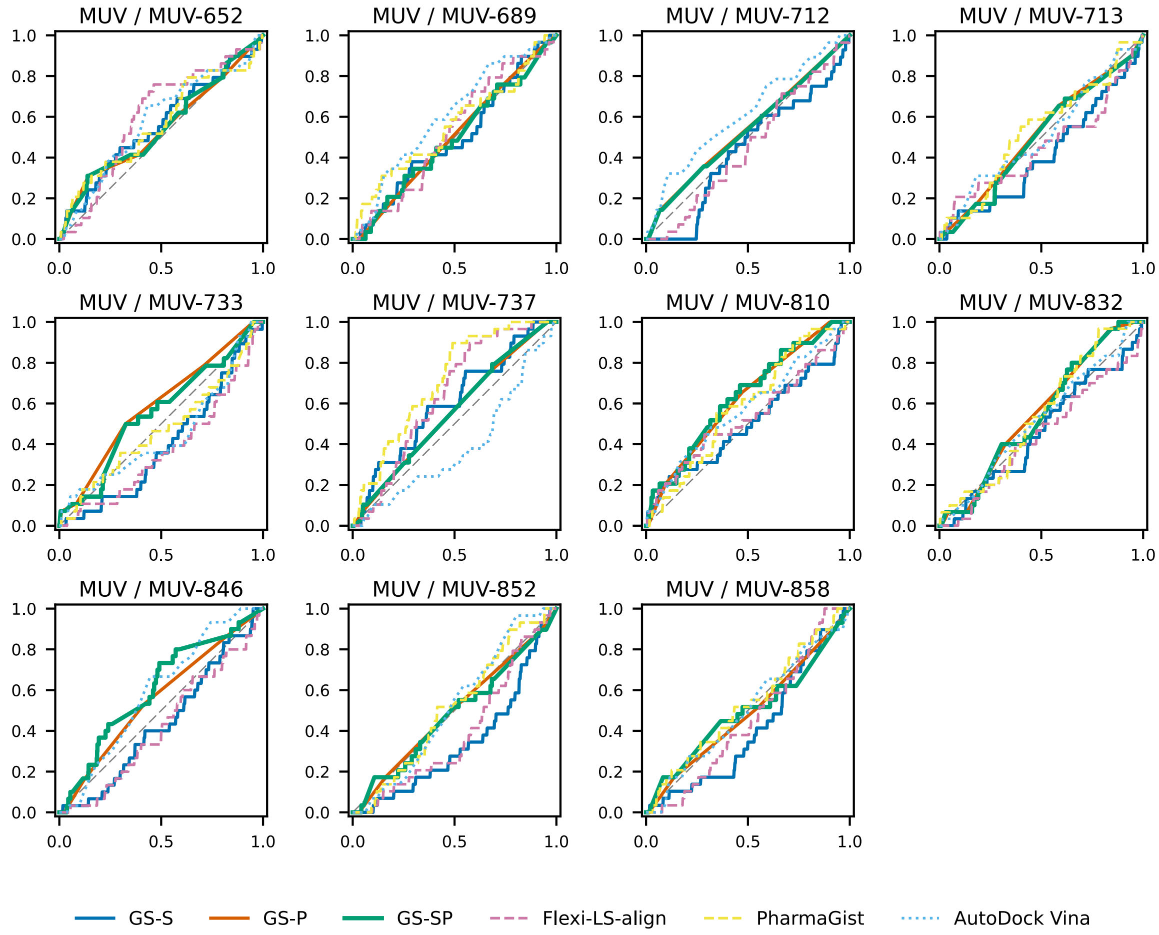

**Figure S1. Full receiver operating characteristic (ROC) curves for all benchmarked targets.** PharmaGist failed to run on two targets, MUV-712 and MUV-846, and was therefore omitted from the corresponding plots.

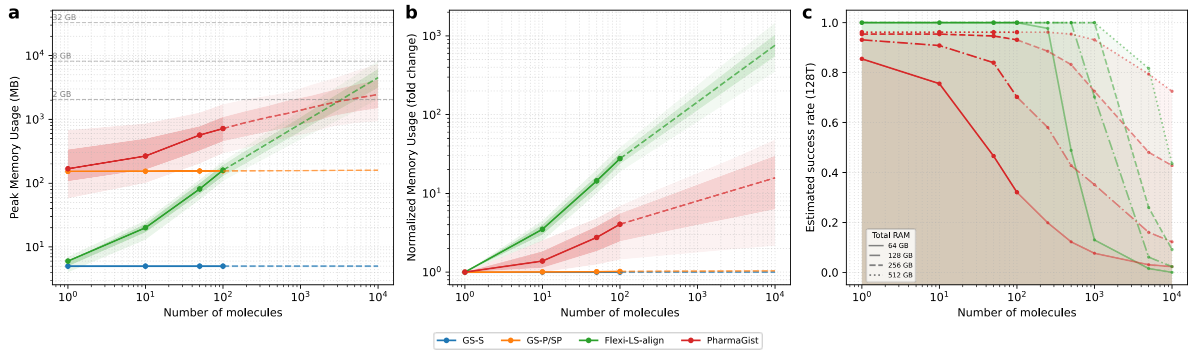

**Figure S2. Empirical memory-scaling behavior of the benchmarked methods across varying workload sizes.**

GS-S and GS-P/SP exhibited near-constant peak memory usage over the tested range, whereas Flexi-LS-align and PharmaGist showed increasing memory usage with workload size and substantially greater target-to-target variability. **(a)** Peak memory usage as a function of the number of molecules processed in a single run. Solid lines denote target-wise medians, with shaded bands indicating the interquartile range and 10th–90th percentile range across targets. Horizontal dashed lines mark representative total RAM budgets (2, 8, and 32 GB). **(b)** Memory usage normalized to the smallest tested workload (1 molecule) for each target and method, highlighting relative growth with increasing workload size. **(c)** Estimated fraction of targets remaining feasible under a 128-thread execution setting for total RAM budgets of 64, 128, 256, and 512 GB, predicted from the observed memory-scaling behavior; GS-S and GS-P/SP are omitted because they showed 100% predicted success across all combinations. In **(a)** and **(b)**, dashed segments indicate extrapolated trends beyond the directly measured workload range; in **(c)**, darker shaded regions indicate extrapolated ranges. As in the main benchmark analysis, PharmaGist failed on two MUV targets; those results were also excluded here.

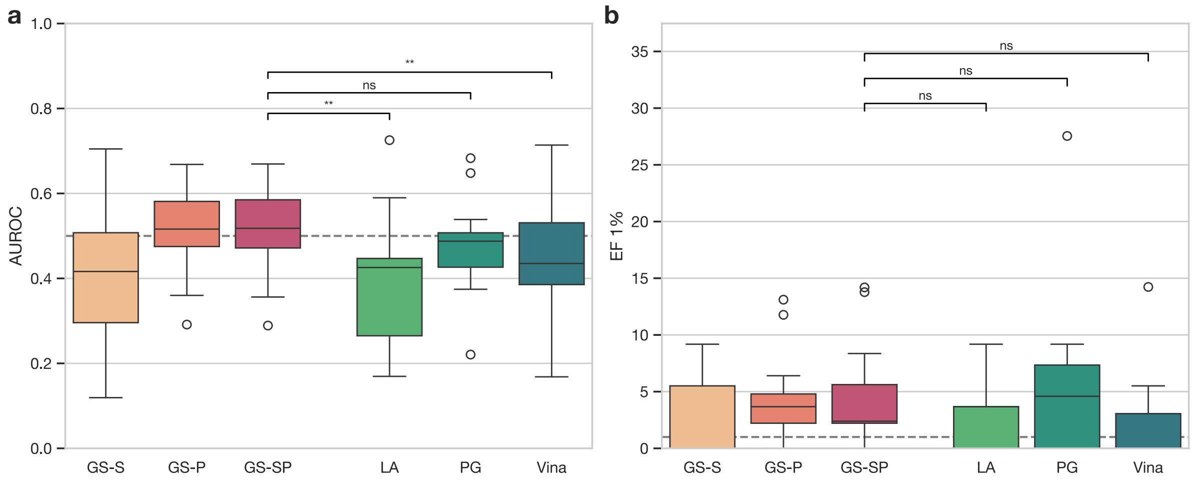

**Figure S3. Performance distributions from the cross-template validation subset of LIT-PCBA.**

Distributions were obtained from clustered reference complexes (up to 5 per target), 10 clustered actives, and 1,000 random decoys are presented, excluding two targets for which only a single reference complex was deposited. Significance brackets indicate pairwise comparisons against GS-SP evaluated using paired Wilcoxon signed-rank tests with Benjamini-Hochberg FDR correction (** $p < 0.01$; ns, not significant). (a) GS-SP showed significantly higher AUROC than Flexi-LS-align and AutoDock Vina, whereas the difference relative to PharmaGist was not significant. (b) No statistically significant pairwise differences were observed for EF1%. LA, Flexi-LS-align; PG, PharmaGist; Vina, AutoDock Vina.

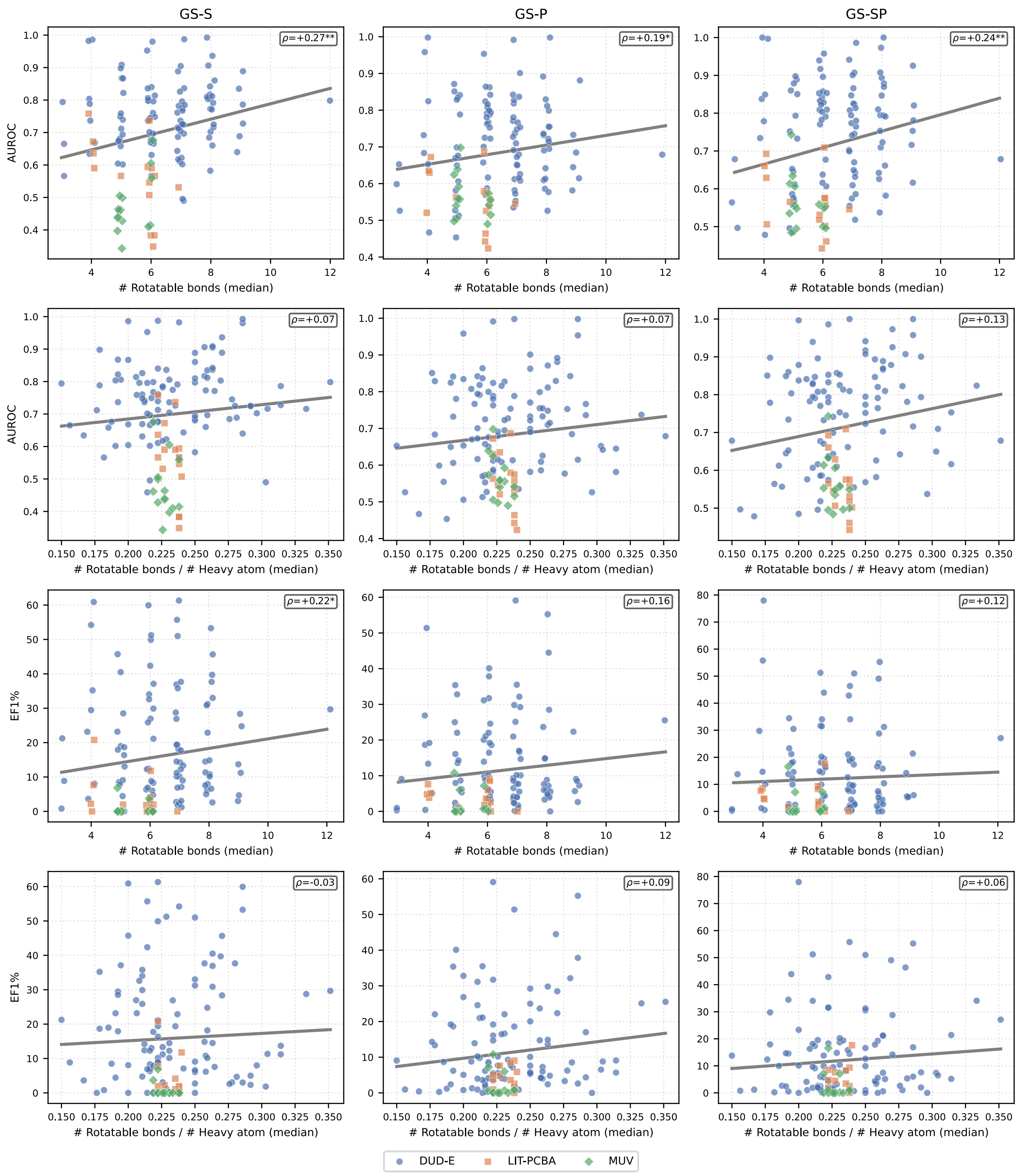

**Figure S4. Target-level relationship between screening performance and ligand flexibility.** Target-level AUROC and EF1% are plotted against ligand flexibility across the DUD-E, LIT-PCBA, and MUV benchmarks for GS-S, GS-P, and GS-SP. Flexibility was quantified using per-target median value of either the raw number of rotatable bonds (first and third rows) or the number of rotatable bonds normalized by heavy-atom count (second and fourth rows). Points denote individual targets and are colored by dataset. Solid lines indicate least-squares trend lines; in-panel annotations report Spearman correlation coefficients (*$p$<0.05; **$p$<0.01). AUROC showed a weak positive association with the raw number of rotatable bonds, whereas this trend became weak or absent after normalization by molecular size. EF1% showed no clear negative association with either flexibility metric, indicating no evidence of a substantial flexibility-dependent loss of performance.

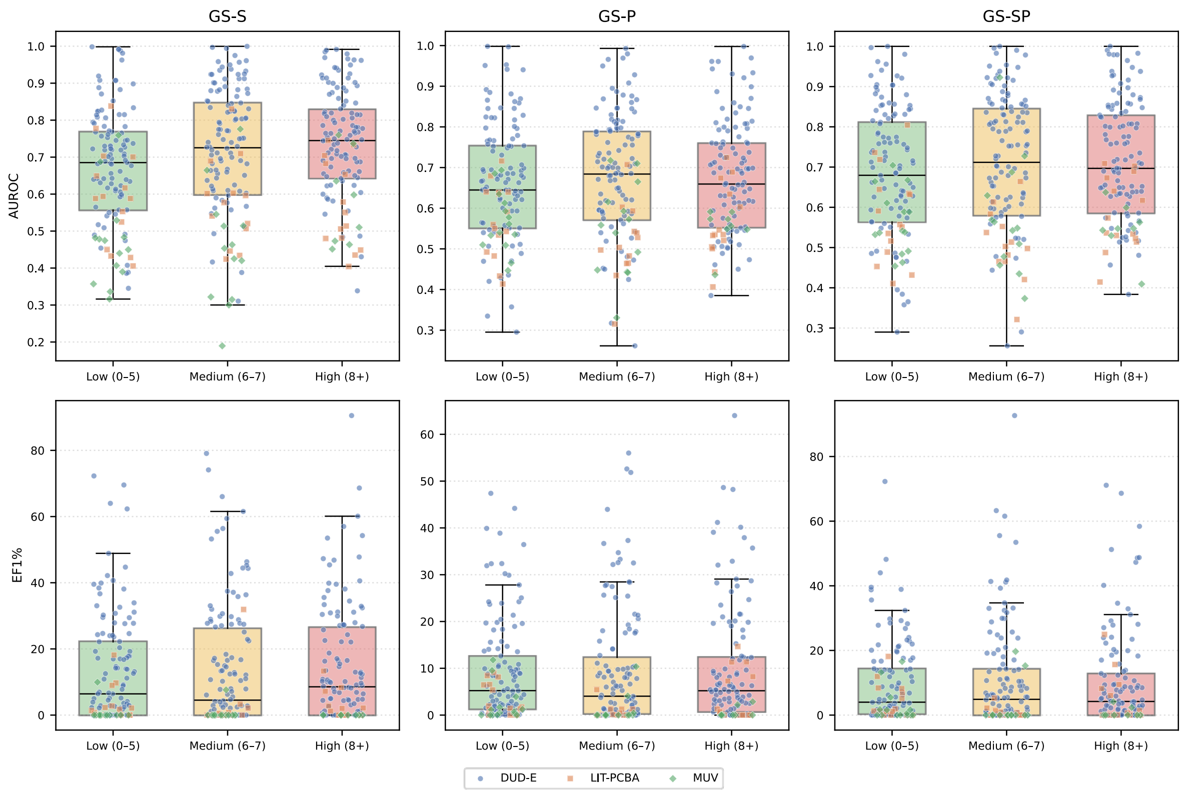

**Figure S5. Stratified analysis of screening performance by ligand flexibility.**
Target-level AUROC and EF1% are shown for low-, medium-, and high-flexibility bins defined by the number of rotatable bonds (low, 0-5; medium, 6-7; high, ≥8), across the DUD-E, LIT-PCBA, and MUV benchmarks. Bin thresholds were chosen to yield approximately equal numbers of molecules per bin across all datasets. Boxplots summarize the target-level distributions for GS-S, GS-P, and GS-SP, with overlaid points representing individual targets colored by dataset. Across all three G-screen variants, no consistent decrease in performance was observed in the highest-flexibility bin, suggesting that ligand flexibility did not impose a strong performance penalty over the range represented in these benchmarks.

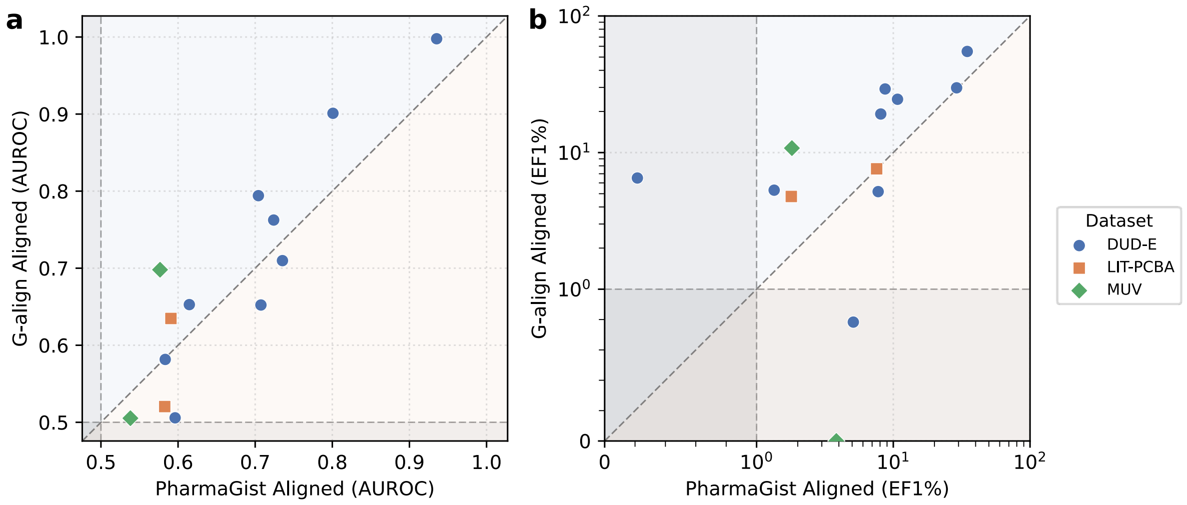

**Figure S6. Alignment-method sensitivity of G-screen scoring.**

Each point corresponds to one target in the subset analysis. (a) AUROC comparison. (b) EF1% comparison, displayed on a symmetric log scale, with the range from 0 to 1 shown linearly. The diagonal indicates equal performance between the two alignment procedures. AUROC was broadly similar between G-align-aligned and PharmaGist-aligned poses, whereas EF1% more often favored G-align-aligned poses, indicating that overall ranking is relatively robust to the alignment method, while early enrichment is more sensitive to the upstream alignment procedure.

### References

(1) Wolber, G.; Langer, T. LigandScout: 3-D Pharmacophores Derived from Protein-Bound Ligands and Their Use as Virtual Screening Filters. *Journal of Chemical Information and Modeling* **2005**, *45* (1), 160–169. DOI: 10.1021/ci049885e.
(2) Abramson, J.; Adler, J.; Dunger, J.; Evans, R.; Green, T.; Pritzel, A.; Ronneberger, O.; Willmore, L.; Ballard, A. J.; Bambrick, J.; et al. Accurate structure prediction of biomolecular interactions with AlphaFold 3. *Nature* **2024**, *630* (8016), 493–500. DOI: 10.1038/s41586-024-07487-w.
(3) Steinegger, M.; Söding, J. MMseqs2 enables sensitive protein sequence searching for the analysis of massive data sets. *Nature Biotechnology* **2017**, *35* (11), 1026–1028. DOI: 10.1038/nbt.3988.
(4) Mirdita, M.; Schütze, K.; Moriwaki, Y.; Heo, L.; Ovchinnikov, S.; Steinegger, M. ColabFold: making protein folding accessible to all. *Nature Methods* **2022**, *19* (6), 679–682. DOI: 10.1038/s41592-022-01488-1.
(5) Consortium, T. U. UniProt: the Universal Protein Knowledgebase in 2025. *Nucleic Acids Research* **2025**, *53* (D1), D609–D617. DOI: 10.1093/nar/gkae1010 (accessed 4/16/2026).
(6) Kim, S.; Chen, J.; Cheng, T.; Gindulyte, A.; He, J.; He, S.; Li, Q.; Shoemaker, Benjamin A.; Thiessen, Paul A.; Yu, B.; et al. PubChem 2025 update. *Nucleic Acids Research* **2025**, *53* (D1), D1516–D1525. DOI: 10.1093/nar/gkae1059 (accessed 4/16/2026).
(7) Goddard, T. D.; Huang, C. C.; Meng, E. C.; Pettersen, E. F.; Couch, G. S.; Morris, J. H.; Ferrin, T. E. UCSF ChimeraX: Meeting modern challenges in visualization and analysis. *Protein Science* **2018**, *27* (1), 14–25. DOI: https://doi.org/10.1002/pro.3235 (accessed 2026/04/15).
(8) Pettersen, E. F.; Goddard, T. D.; Huang, C. C.; Meng, E. C.; Couch, G. S.; Croll, T. I.; Morris, J. H.; Ferrin, T. E. UCSF ChimeraX: Structure visualization for researchers, educators, and developers. *Protein Science* **2021**, *30* (1), 70–82. DOI: https://doi.org/10.1002/pro.3943 (accessed 2026/04/15).
(9) Meng, E. C.; Goddard, T. D.; Pettersen, E. F.; Couch, G. S.; Pearson, Z. J.; Morris, J. H.; Ferrin, T. E. UCSF ChimeraX: Tools for structure building and analysis. *Protein Science* **2023**, *32* (11), e4792. DOI: https://doi.org/10.1002/pro.4792 (accessed 2026/04/15).
